# Optimal maturation of the SIV-specific CD8^+^ T-cell response after primary infection is associated with natural control of SIV. ANRS SIC study

**DOI:** 10.1101/2019.12.20.885459

**Authors:** Caroline Passaes, Antoine Millet, Vincent Madelain, Valérie Monceaux, Annie David, Pierre Versmisse, Naya Sylla, Emma Gostick, David A. Price, Antoine Blancher, Nathalie Dereuddre-Bosquet, Gianfranco Pancino, Roger Le Grand, Olivier Lambotte, Michaela Müller-Trutwin, Christine Rouzioux, Jeremie Guedj, Veronique Avettand-Fenoel, Bruno Vaslin, Asier Sáez-Cirión

**Author notes:** These authors contributed equally to this work (Bruno Vaslin, Asier Sáez-Cirión).

## Abstract

Highly efficient virus-specific CD8^+^ T-cells are associated with immune control of HIV infection, but it remains unclear how these cells are generated and maintained over time. We used a macaque model of spontaneous control of SIVmac251 infection to monitor the development and evolution of potent antiviral CD8^+^ T-cell responses. SIV-specific CD8^+^ T-cells emerged during primary infection in all animals. However, the ability of CD8^+^ T cells to suppress SIV replication was low in early stages but increased after a period of maturation, temporally linked with the establishment of sustained low-level viremia in controller macaques. SIV-specific CD8^+^ T-cells with a central memory phenotype expressed higher levels of survival markers in controllers *versus* non-controllers. In contrast, a persistently skewed differentiation phenotype was observed among central memory SIV-specific CD8^+^ T-cells in non-controllers since primary infection, typified by relatively high expression levels of T-bet.

Collectively, these data show that the phenotype of SIV-specific CD8^+^ T-cells defined early after SIV infection favor the gain of antiviral potency as a function of time in controllers, whereas SIV-specific CD8^+^ T-cell responses in non-controllers fail to gain antiviral potency due to early defects imprinted in the central memory pool.

## INTRODUCTION

The ability of CD8^+^ T-cells to control viral replication has been extensively documented in the setting of HIV/SIV infection (McBrien et al., 2018; Walker and McMichael, 2012). Primary infection is characterized by massive viremia, which subsides following the expansion of HIV/SIV-specific CD8^+^ T-cells (Borrow et al., 1994; Koup et al., 1994). However, the virus is not eradicated, leading to the emergence of immune escape variants (Allen et al., 2000; Borrow et al., 1997; O’Connor et al., 2002; Price et al., 1997) and to CD8^+^ T-cell exhaustion during the chronic phase of infection (Day et al., 2006; Petrovas et al., 2006; Petrovas et al., 2007; Trautmann et al., 2006). These observations suggest that naturally generated HIV/SIV-specific CD8^+^ T-cells are frequently suboptimal in terms of antiviral efficacy, potentially reflecting limited cross-reactivity and/or intrinsic defects in the arsenal of effector functions required to eliminate infected CD4^+^ T-cells (Du et al., 2016; Lecuroux et al., 2013). The latter possibility is especially intriguing in light of *ex vivo* experiments showing that effective suppression of viral replication is a particular feature of CD8^+^ T-cells isolated from HIV controllers (HICs) (Angin et al., 2016; Saez-Cirion et al., 2007; Saez-Cirion et al., 2009; Tansiri et al., 2015).

HICs are a rare group of individuals who control viremia to very low levels without antiretroviral therapy (Saez-Cirion and Pancino, 2013). Understanding the mechanisms associated with such spontaneous control of HIV infection seems crucial for the development of new strategies designed to achieve remission. Efficient CD8^+^ T-cell responses are almost universally present in HICs (Betts et al., 2006; Chowdhury et al., 2015; Hersperger et al., 2011a; Hersperger et al., 2010; Migueles et al., 2002; Migueles et al., 2008; Saez-Cirion et al., 2007; Saez-Cirion et al., 2009; Zimmerli et al., 2005). These individuals also frequently express the protective human leukocyte antigen (HLA) allotypes HLA-B*27 and HLA-B*57, further supporting a key role for CD8^+^ T-cells in the natural control of HIV (Lecuroux et al., 2014; Migueles et al., 2000; Pereyra et al., 2008). However, the presence of protective HLA alleles is neither sufficient nor necessary for natural control of infection, and HICs carrying non-protective HLA class I alleles also carry CD8^+^ T-cells with strong HIV suppressive capacity (Lecuroux et al., 2014). Although the qualitative properties of CD8^+^ T-cells from HICs have been extensively characterized, these analyses have been essentially performed during chronic infection, when viremia was already under control, often several years after the acquisition of HIV. It therefore remains unclear how these high-quality CD8^+^ T-cell responses develop from the early stages of infection and evolve over time.

Cynomolgus macaques (CyMs, *Macaca fascicularis*) infected with SIVmac251 closely recapitulate the dynamics and key features of HIV infection, including similar levels of viral replication in the acute and chronic phases of infection, memory CD4^+^ T-cell depletion, rapid seeding of the viral reservoir, and eventually progression to AIDS with diarrhea, weight loss, high incidence of lymphoblastic lymphomas and marked decrease of CD4+ T cells within 145 to 464 days post-infection (Antony and MacDonald, 2015; Feichtinger et al., 1990; Karlsson et al., 2007; Mannioui et al., 2009; Putkonen et al., 1989). As in humans, some individuals control infection naturally in the absence of treatment. CyMs from Mauritius offer the additional advantage of limited MHC diversity, making them particularly attractive for the study of CD8+ T-cell responses. Indeed, natural SIV control in Mauritius CyMs is favored by the presence of the MHC haplotype M6 (Aarnink et al., 2011; Mee et al., 2009). Natural SIV control can be also achieved in CyMs inoculated with a relatively low virus dose exposure through the intra rectal route (*i.r.*), independent of their MHC haplotype (Bruel et al., 2015).

We therefore took advantage of these validated CyM models spreading from natural SIV control to progression to AIDS to study the dynamics of SIV-specific CD8^+^ T-cell responses in blood and tissues from the onset of infection in both SIV controllers and viremic macaques. Using this approach, we identified an optimal maturation pathway that enabled SIV-specific CD8^+^ T-cells to acquire potent antiviral functions, control viremia, and survive in SICs.

## RESULTS

### SIV controllers are characterized by partial restoration of CD4+ T-cell counts and progressive decline in the frequency of SIV-carrying cells in blood

We monitored prospectively the outcome of infection in 12 SIV controllers (SICs) and 4 viremic CyMs (VIRs) inoculated *i.r.* with SIVmac251. These animals carried or not the protective M6 haplotype and were inoculated with 5 or 50 animal infectious dose_50_ (AID_50_) of SIVmac251 (Supplemental Table 1). SIV controllers decreased plasma viral load (VL) to levels below 400 SIV-RNA copies/mL, at least twice, over a follow up period of 18 months, while VIRs consistently maintained VL above 400 SIV-RNA copies/mL. The threshold of 400 RNA copies/mL was chosen in coherence with our studies in human cohorts of natural HIV control (Angin et al., 2016; Noel et al., 2016; Saez-Cirion et al., 2013; Saez-Cirion et al., 2007; Saez-Cirion et al., 2009). Ten SICs achieved control of viremia within 3 months. The other two SICs (BL669 and BO413) achieved VL below 400 SIV-RNA copies/mL for the first time 14 months after inoculation. One VIR CyM (AV979) developed a tonsilar lymphoma, an AIDS related event reported at high frequency in this species upon SIV infection (Feichtinger et al., 1990).

Some differences in peak viremia were observed between SICs and VIRs (Figure 1A, Table 1). These differences became more pronounced over time (Figure 1A), because plasma viremia was suppressed more rapidly in SICs versus VIRs (Table 1). Levels of cell-associated SIV-DNA in blood from SICs and VIRs were comparable before peak viremia, but differences became apparent as plasma VLs declined and were maintained throughout chronic infection (Figure 1B, Table 1). In addition, CD4^+^ T-cell counts declined markedly in blood from both SICs and VIRs during primary infection (Figure 1C, Table 1). Subsequently, a degree of recovery was observed in SICs, whereas further gradual decline was observed in VIRs (Figure 1C, Table 1).

**Figure 1.**
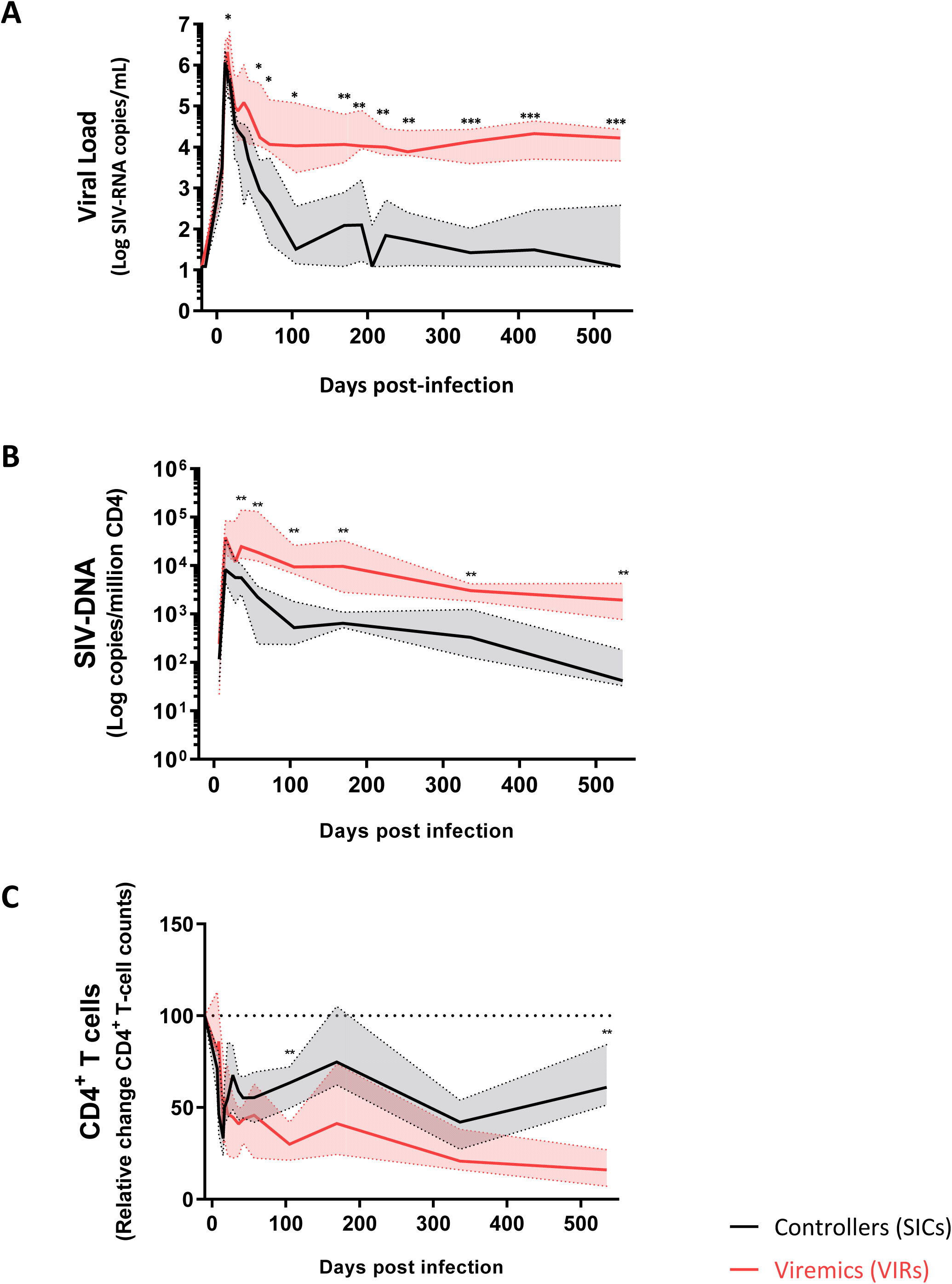
SIV controllers are characterized by partial restoration of CD4^+^ T-cell counts and progressive decline in the frequency of SIV-carrying cells in blood. (**A**) Plasma VL kinetics, (**B**) kinetics of SIV-DNA levels in blood and (**C**) longitudinal evolution of CD4^+^ T-cell counts (results are shown as fold-change in absolute CD4^+^ T-cell counts relative to baseline) in blood in SIV controllers (SICs, black) and viremic CyMs (VIRs, red). Median and interquartile range are shown. *p < 0.05, **p < 0.01, ***p < 0.001; Mann-Whitney U-test.

**Table 1.** Virologic and immunologic characteristics from SICs *versus* VIRs.

|  | Controllers | Viremics | p |
| --- | --- | --- | --- |
| **RNA viral load** |  |  |  |
| Peak (Log SIV-RNA copies/mL) | 6.1<br>[5.1 – 7.1] | 6.6<br>[6.3 – 6.9] | 0.058 |
| Time to peak (days *p.i.*) | 12.5<br>[11 – 17] | 14<br>[14 – 17] | 0.239 |
| Set-point<sup>#</sup> (Log SIV-RNA copies/mL) | 1.5<br>[1.1 – 3.6] | 4.0<br>[3.2 – 5.2] | **0.014** |
| Slope after peak viremia (1/slope peak – set-point) | $-1.3 \times 10^{-4}$<br>[$-1.5 \times 10^{-3}$ to $-2.1 \times 10^{-5}$] | $-5.5 \times 10^{-5}$<br>[$-1.2 \times 10^{-4}$ to $-2.3 \times 10^{-5}$] | 0.058 |
| **DNA viral load** |  |  |  |
| Peak (Log SIV-DNA copies/million CD4) | 4.0<br>[3.5 – 4.7] | 4.6<br>[4.1 – 5.3] | 0.133 |
| Time to peak (days *p.i.*) | 15<br>[15 – 36] | 15<br>[15 – 36] | 1 |
| Set-point<sup>#</sup> (Log SIV-DNA copies/million CD4) | 2.7<br>[2.1 – 3.5] | 4.0<br>[3.8 – 4.5] | **0.002** |
| Descending slope (1/slope peak – set-point) | $-9.0 \times 10^{-3}$<br>[$-4.8 \times 10^{-2}$ to $-1.0 \times 10^{-3}$] | $-4.9 \times 10^{-3}$<br>[$-1.2 \times 10^{-2}$ to $-4.4 \times 10^{-4}$] | 0.262 |
| **CD4<sup>+</sup> T-cell counts** |  |  |  |
| Nadir CD4<sup>+</sup> T-cells (cells/μL blood) | 238<br>[71 – 910] | 179<br>[73 – 276] | 0.379 |
| Time to nadir CD4<sup>+</sup> T-cells (days) | 15<br>[9 – 36] | 19<br>[15 – 28] | 0.122 |
| Set-point<sup>#</sup> CD4<sup>+</sup> T-cells (cells/μL blood) | 602<br>[257 – 1468] | 154<br>[80 – 363] | **0.008** |

**Median and range are indicated. p, Mann-Whitney U-test. #Set point defined as Month 3 post-infection**

These results evidenced the distinctive dynamics of SIV infection in SICs and VIRs, characterized by very modest differences during the early weeks following inoculation that were progressively exacerbated during transition to chronic infection. The differences between SICs and VIRs during the chronic phase of SIV infection were consistent with the observations in human cohorts of HIV controllers.

### SIV control is associated with early preservation of lymph nodes

To characterize the extent of SIV control in greater depth, we monitored CD4^+^ T-cells and total SIV-DNA longitudinally in peripheral lymph nodes (PLNs) and rectal biopsies (RBs). At the end of the study, we conducted similar evaluations in bone marrow, spleen, mesenteric lymph nodes (MLNs), and colonic mucosa, comparing SICs versus VIRs. The frequency of CD4^+^ T-cells similarly declined in RBs from both SICs and VIRs during the acute stage of primary infection. While the frequency of CD4^+^ T-cells was later partially restored in SICs, it continued to decline in VIRs (Figure 2A). These results matched the observations in blood samples (Figure 1B). In contrast, the frequency of CD4^+^ T-cells was maintained close to baseline in PLNs from SICs, even during primary infection (day 14 post-infection [*p.i.]*), but steadily declined over time in VIRs (Figure 2B). At the time of euthanasia, CD4^+^ T-cell frequencies were substantially higher in blood (Figure 1C), bone marrow, spleen, PLNs, MLNs, and colonic mucosa (Figure 2C) from SICs versus VIRs.

**Figure 2.**
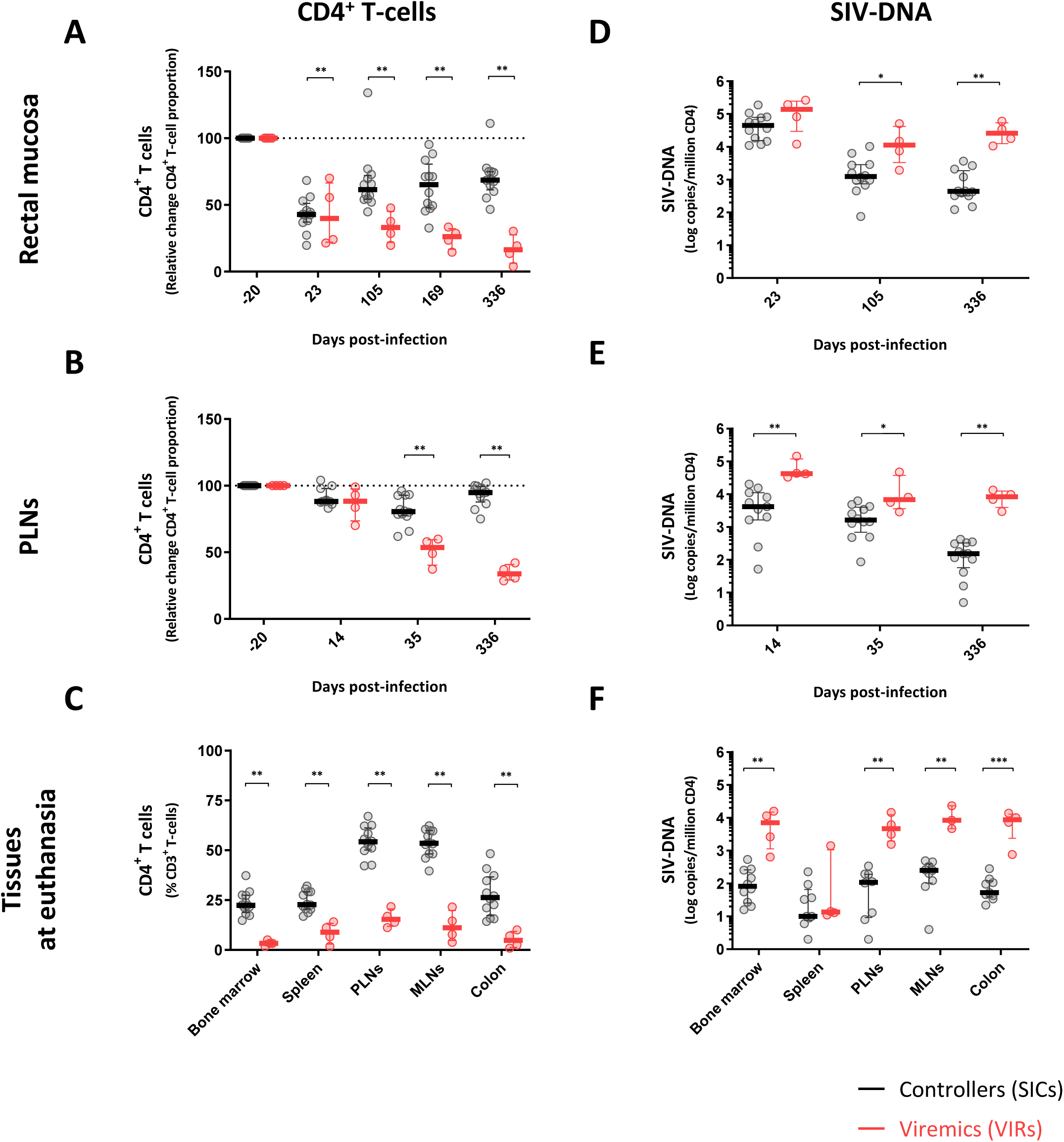
SIV control is associated with early preservation of lymph nodes. (**A**–**B**) Longitudinal evolution of CD4^+^ T-cells in rectal mucosa (**A**), and peripheral lymph nodes (**B**) in SIV controllers (SICs, black) and viremic CyMs (VIRs, red). Results in rectal mucosa and peripheral lymph nodes are shown as fold-change in percent frequencies of CD3^+^ CD4^+^ T-cells among CD3^+^ lymphocytes relative to baseline. (**C**) Percent frequencies of CD3^+^ CD4^+^ T-cells among CD3^+^ lymphocytes in bone marrow, spleen, peripheral and mesenteric lymph nodes, and colon mucosa at euthanasia. (**D**–**E**) Kinetics of SIV-DNA levels in rectal mucosa (**D**), and peripheral lymph nodes (**E**) in SIV controllers (black) and viremic CyMs (red). (**F**) Levels of SIV-DNA in bone marrow, spleen, peripheral and mesenteric lymph nodes, and colon at euthanasia. Results are expressed as copies SIV-DNA/million CD4^+^ T-cells. Median and interquartile range are shown. *p < 0.05, **p < 0.01; Mann-Whitney U-test.

Cell-associated SIV-DNA levels closely mirrored the dynamics of CD4^+^ T-cells. Similarly high levels of cell-associated SIV-DNA were observed in RBs from SICs and VIRs during primary infection, but lower levels were observed in RBs from SICs *versus* VIRs during chronic infection (Figure 2D). Of note, SIV-DNA levels were already approximately 1 log lower in PLNs *versus* RBs from SICs during primary infection, and accordingly, lower levels were observed in PLNs from SICs *versus* VIRs since day 14 *p.i.* (Figure 2E). This finding suggests that early viral replication may be contained more efficiently in lymphoid nodes in SICs compared with other explored anatomical compartments. Moreover, SIV-DNA was also detected in alveolar macrophages from all CyMs throughout the course of infection, again at lower levels in SICs *versus* VIRs during chronic infection (Figure S1A, bottom panel). In addition, SIV-DNA levels trended to decline progressively over time in SICs in all tissues analyzed, whereas SIV-DNA levels remained stable after primary infection in VIRs (Figure 1B, 2D-F). At the time of euthanasia, SIV-DNA levels were substantially lower in blood (Figure 1B), bone marrow, PLNs, MLNs, and gut mucosa (Figure 2F and Figure S1B) from SICs *versus* VIRs.

Collectively, these data indicate that progressive systemic control of viral replication is achieved in SICs with CD4^+^ T-cell preservation and lower pan-anatomical reservoirs of SIV-DNA. Our results also underline the early preservation of PLNs in these animals.

### The dynamics of CD8^+^ T-cells expansion and activation do not predict control of SIV

To understand the mechanisms that contribute to immune control of SIV, we first monitored the proliferation and activation dynamics of total CD8^+^ T-cells in blood and lymphoid tissues from SICs and VIRs. Recent studies in cohorts of hyperacute HIV infected individuals indicate that the changes observed in the total CD8^+^ T-cell activation during acute infection may be largely related to changes in the HIV-specific CD8^+^ T-cell pool (Ndhlovu et al., 2015; Takata et al., 2017). The frequencies of CD8^+^ T-cells expressing Ki-67 in blood increased to maximum levels during primary infection (measured peak at day 15 *p.i.*), coinciding with the measured peak of viremia, then declined steadily to baseline levels during chronic infection (Figure 3A). Similar dynamics were observed in PLNs (Figure 3B) and gut mucosa (Figure 3C). In general, there were no significant differences between SICs and VIRs with respect to the dynamics of Ki-67 expression within the CD8^+^ T-cell pool, although lower frequencies of CD8^+^ T-cells expressing Ki-67 were observed during chronic infection in PLNs from SICs *versus* VIRs (Figure 3B).

**Figure 3.**
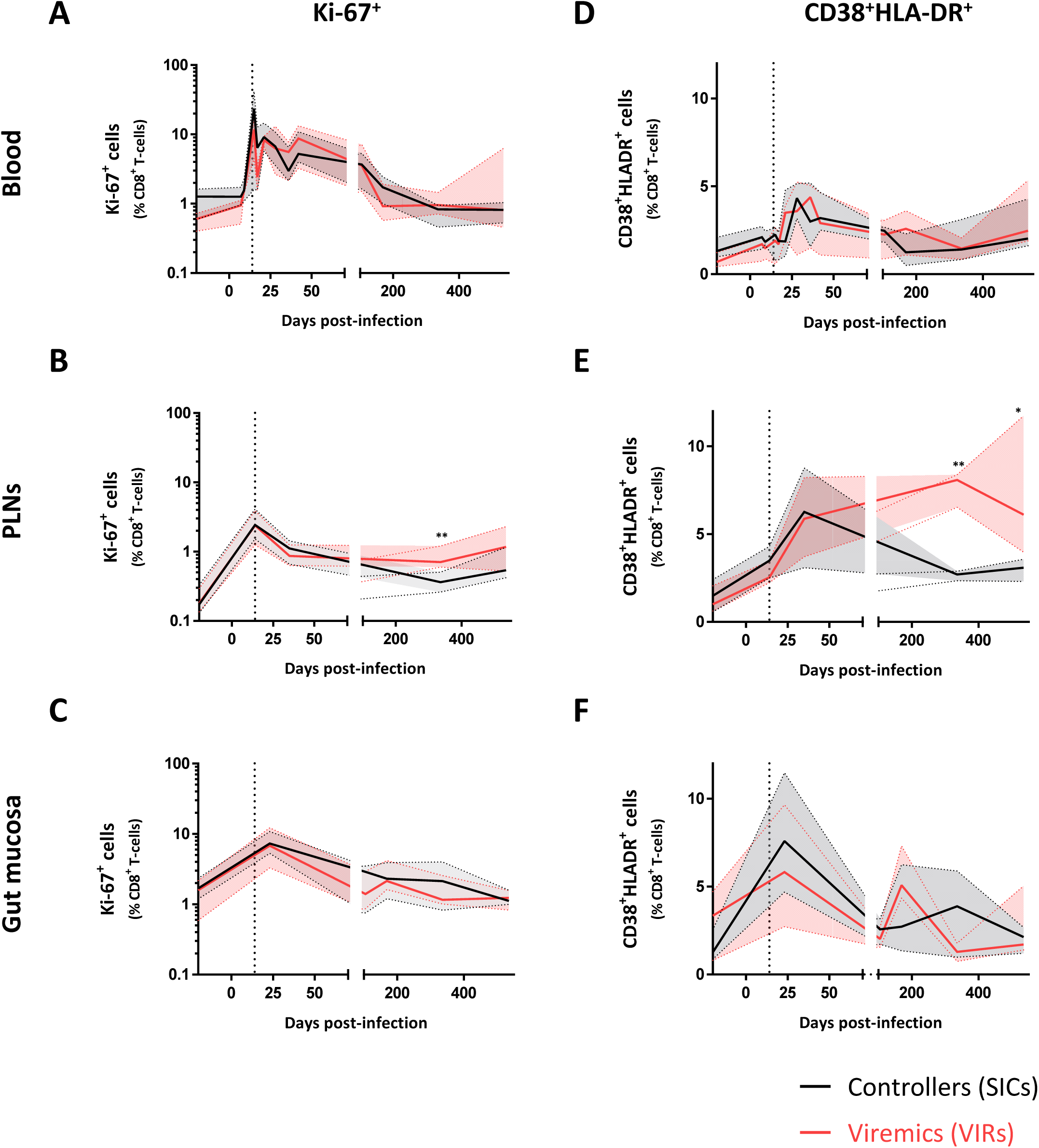
The dynamics of CD8^+^ T-cells expansion and activation do not predict control of SIV. (**A**–**C**) Evolution of Ki-67^+^ CD8^+^ T-cells in blood (**A**), peripheral lymph nodes (**B**), and rectal mucosa (**C**) in SIV controllers (black) and viremic CyMs (red). (**D**–**F**) Evolution of CD38^+^ HLA-DR^+^ CD8^+^ T-cells in blood (**D**), peripheral lymph nodes (**E**), and rectal mucosa (**F**) in SIV controllers (black) and viremic CyMs (red). Median and interquartile range are shown. Vertical dashed lines indicate peak VLs. *p < 0.05, **p < 0.01; Mann-Whitney U-test.

The frequencies of CD8^+^ T-cells expressing the activation markers CD38 and HLA-DR in blood, PLNs, and gut mucosa increased similarly during primary infection (measured peak at day 28 *p.i.*), following the dynamics of Ki-67 expression in the same compartments (Figure 3D–F). Again, there were no significant differences between SICs and VIRs with respect to the early dynamics of total CD8^+^ T-cell activation, but lower frequencies of CD8^+^ T-cells expressing CD38 and HLA-DR were observed during chronic infection in PLNs from SICs *versus* VIRs (Figure 3E).

Overall, our findings indicate that although lower activation and proliferation is observed in of CD8^+^ T-cells from SICs than VIRs in the chronic stage of infection, the early proliferation and activation dynamics of the total pool of CD8^+^ T-cells do not distinguish subsequent progression rates.

### SIV-specific CD8^+^ T-cell frequencies do not predict control of SIV

In parallel experiments, we analyzed CD8^+^ T-cell responses to a pool of optimal SIVmac251 peptides, which included peptides from different SIV proteins recognized by the most frequent MHC haplotypes in CyMs (M1, M2 and M3) and by the MHC haplotype M6 (Supplemental Table 2). All the animals carried at least one haplotype matching some peptide, and overall there was not difference in the number of peptides tested theoretically recognized in controllers and non-controllers (p=0.35). SIV-specific CD8^+^ T-cells producing TNFα (cytokine showing the lowest background in the absence of peptide or in presence of peptide during the baseline and hence used as reference) emerged in all CyMs during primary infection, coinciding with the peak of viremia, and no significant differences were observed between SICs and VIRs with respect to the frequencies of these cells in any anatomical compartment at any stage of infection (Figure 4A). Similarly, no consistent differences were observed between SICs and VIRs with respect to the frequencies of SIV-specific CD8^+^ T-cells that produced other cytokines, including IFNγ (Figure S2A and S3A) and IL-2 (Figure S2B and S3B), or mobilized CD107a (Figure S2C and S3C). The overall SIV-specific CD8^+^ T-cell response, determined in each CyM as the frequency of cells displaying at least one function (TNFα, IFNγ, IL-2, or CD107a), was also equivalent between SICs and VIRs across anatomical compartments during primary and chronic infection (Figure S2D and S3D). In addition, no clear differences between SICs and VIRs were observed with respect to the frequencies of SIV-specific CD8^+^ T-cells displaying at least three functions simultaneously in blood or PLNs during acute infection, but higher frequencies of polyfunctional SIV-specific CD8^+^ T-cells were present during chronic infection in lymphoid tissues from SICs *versus* VIRs (Figure 4B and S4). Of note, no differences in the magnitude (Figure S5A) and polyfunction (Figure S5B) of SIV-specific CD8^+^ T-cells from SICs and VIRs were observed either when a pool of overlapping peptides spanning SIV Gag was used instead of the optimal peptide pool to stimulate the cells.

**Figure 4.**
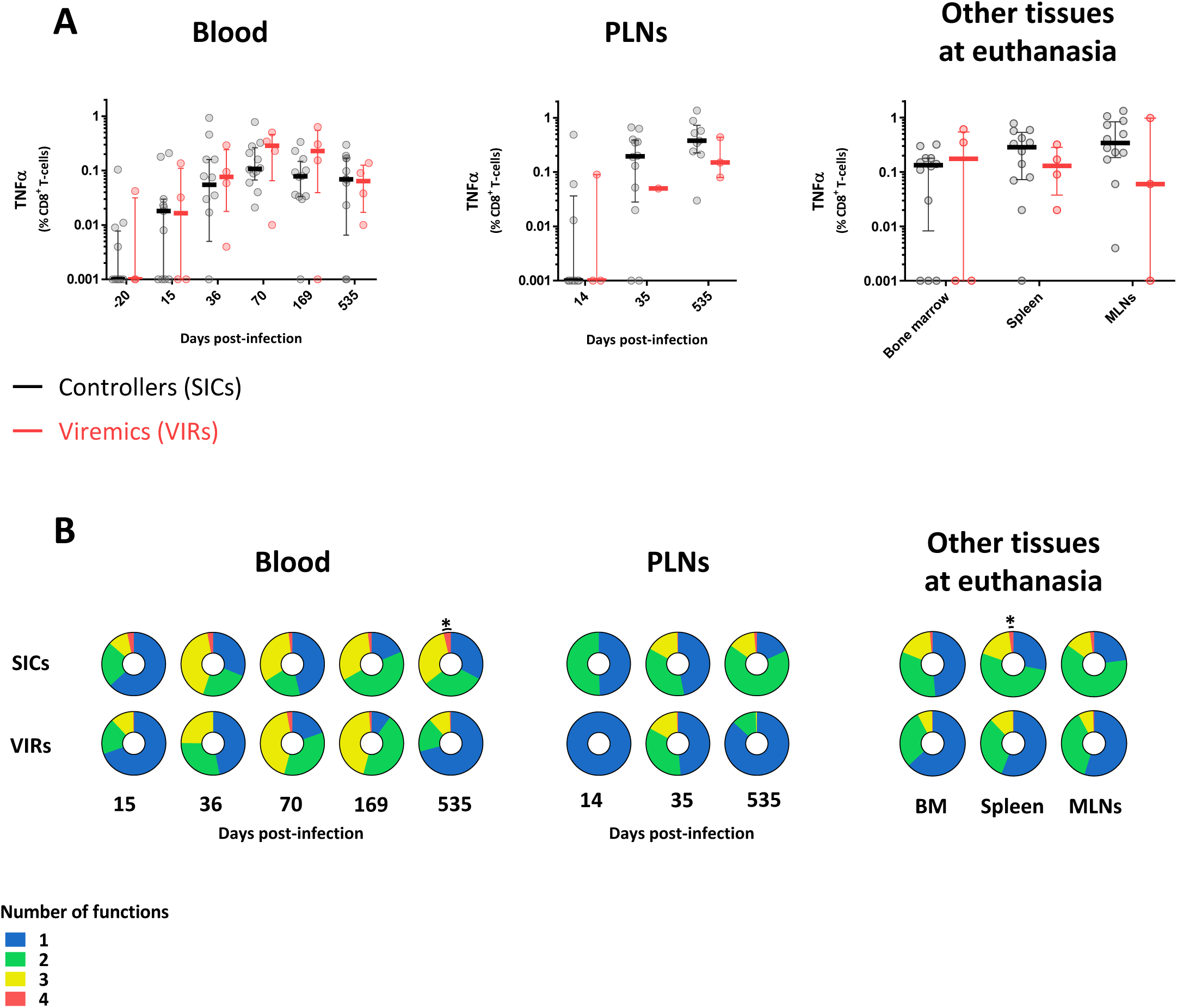
SIV-specific CD8^+^ T-cell frequencies do not predict control of SIV. (**A**) TNFα production by SIV-specific CD8^+^ T-cells in blood and peripheral lymph nodes over the course of infection and in bone marrow, spleen, and mesenteric lymph nodes at euthanasia in SIV controllers (black) and viremic CyMs (red). Results are shown as percent frequencies among CD8^+^ T-cells. Median and interquartile range are shown. (**B**) Functional profiles of SIV-specific CD8^+^ T-cells in blood and peripheral lymph nodes over the course of infection and in bone marrow, spleen, and mesenteric lymph nodes at euthanasia in SIV controllers (black) and viremic CyMs (red). Doughnut charts show median percent frequencies of SIV-specific CD8^+^ T-cells expressing IFNγ, TNFα, IL-2, and/or CD107a. Colors indicate number of simultaneous functions (blue, 1; green, 2; yellow, 3; red, 4). *p < 0.05; Mann-Whitney U-test.

These data suggest that natural control of SIV is not associated with acutely generated, functionally superior SIV-specific CD8^+^ T-cell responses, defined on the basis of cytokine production and degranulation.

### Progressive acquisition of CD8^+^ T-cell-mediated SIV-suppressive activity is associated with control of SIV

CD8^+^ T-cells from HICs typically suppress *ex vivo* infection of autologous CD4^+^ T-cells (Angin et al., 2016; Buckheit et al., 2012; Julg et al., 2010; Saez-Cirion et al., 2007; Saez-Cirion et al., 2009; Tansiri et al., 2015). We therefore investigated this property as a potential discriminant between SICs and VIRs. The capacity of CD8^+^ T-cells in blood and PLNs to suppress infection of autologous CD4^+^ T-cells was relatively weak in all CyMs during acute infection (Figure 5A), but remarkably, this activity correlated negatively with viremia on day 15 *p.i.* (Figure 5B, upper panel) suggesting its contribution to control viremia since early time points. Interestingly, the CD8^+^ T-cell-mediated SIV-suppressive activity increased substantially over time in SICs (Figure 5A and S6), either in blood or tissues. No such acquisition of SIV-suppressive activity was observed in VIRs (Figure 5A and S6). Moreover, CD8^+^ T-cell-mediated SIV-suppressive activity on day 70 *p.i.* correlated negatively (or trended to correlate) with all subsequent determinations of plasma VL (Figure S7) and there was a negative correlation between the CD8^+^ T-cell-mediated SIV-suppressive activity at euthanasia and the viremia at this time (Figure 5C, upper panel). In contrast, no significant correlations were identified at any time point between SIV-specific CD8^+^ T-cell frequencies, categorized according to TNFα production in response to SIV peptides, and measurements of plasma VL (Figure 5B-C, bottom panels). Moreover, CD8^+^ T-cell-mediated SIV-suppressive activity across the entire follow-up period, quantified as area under the curve, trended to correlate negatively with plasma VL (r_s_ = –0.47, p = 0.07), whereas no such association was identified for the frequency of SIV-specific CD8^+^ T-cells (r_s_ = –0.01, p = 0.97) (Figure 5D).

**Figure 5.**
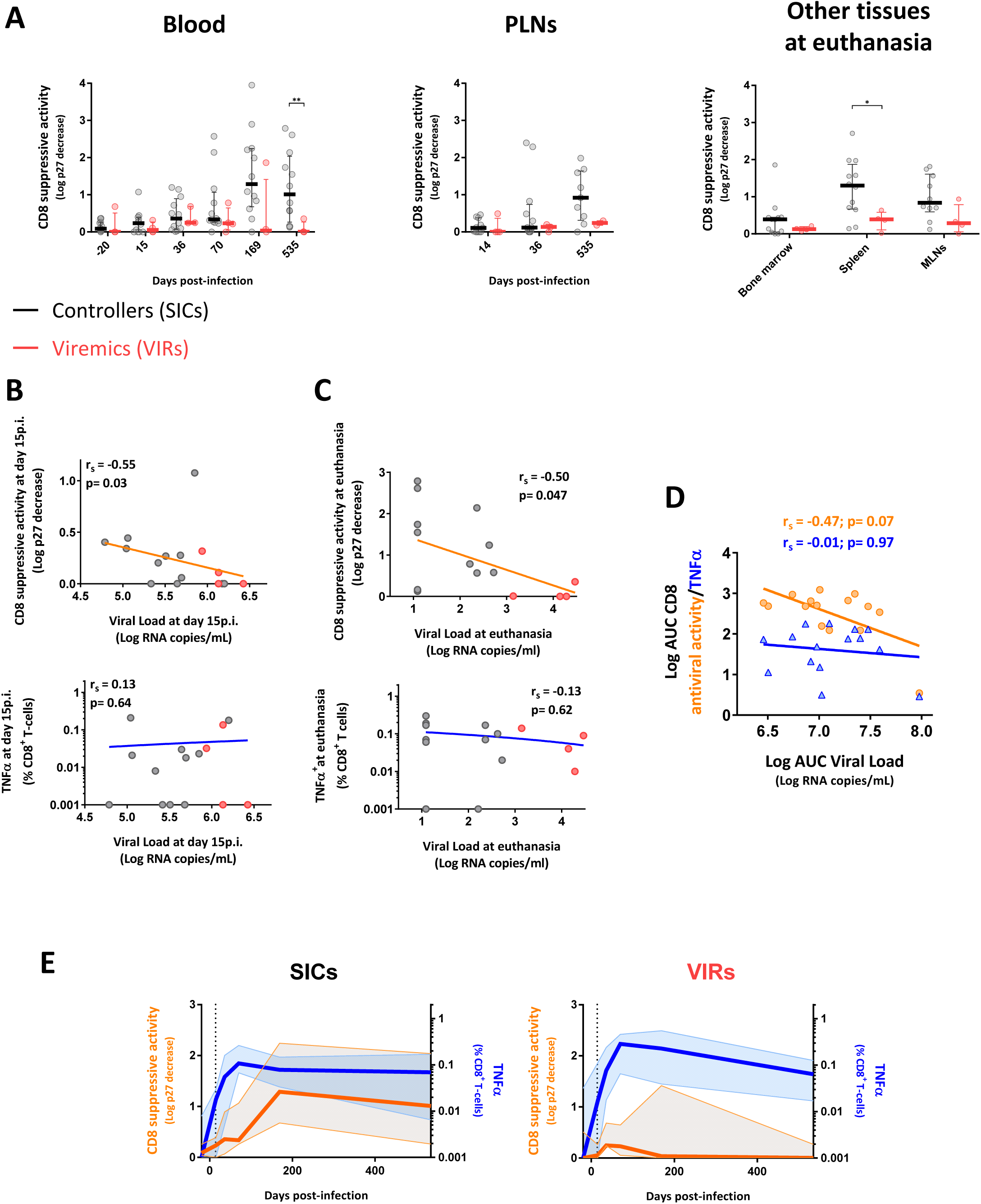
Progressive acquisition of CD8^+^ T-cell-mediated SIV-suppressive activity is associated with control of SIV. (**A**) CD8^+^ T-cell-mediated SIV-suppressive activity in blood and peripheral lymph nodes over the course of infection and in bone marrow, spleen, and mesenteric lymph nodes at euthanasia in SIV controllers (black) and viremic CyMs (red). Results are shown as log p27 decrease in the presence of CD8^+^ T-cells. *p < 0.05, **p < 0.01; Mann-Whitney U-test. (**B**) Spearman correlations between CD8^+^ T-cell-mediated SIV-suppressive activity (upper panel) or TNFα production by SIV-specific CD8^+^ T-cells (bottom panel) on day 15 *p.i.* with plasma VL on day 15 *p.i.* Grey symbols, SIV controllers; red symbols, viremic CyMs. (**C**) Spearman correlations between CD8^+^ T-cell-mediated SIV-suppressive activity (upper panel) or TNFα production by SIV-specific CD8^+^ T-cells (bottom panel) at euthanasia with plasma VL at euthanasia. Grey symbols, SIV controllers; red symbols, viremic CyMs (**D**) Spearman correlations between area under the curve (AUC) for plasma VL and AUC for CD8^+^ T-cell-mediated SIV-suppressive activity (orange) and between AUC for plasma VL and AUC for TNFα production by SIV-specific CD8^+^ T-cells (blue). AUC for plasma VL, TNFα production and CD8+ T-cell antiviral activities were calculated using the sequential values obtained throughout the duration of our study in the blood of the infected animals (Figure S6). (**E**) Side by side comparison of the longitudinal kinetics of TNFα production by SIV-specific CD8^+^ T-cells in blood shown in figure 4A (blue) and CD8^+^ T-cell-mediated SIV-suppressive activity in blood shown in figure 5A(orange) in SIV controllers (left panel) and viremic CyMs (right panel). Median and interquartile range are shown.

In a complementary study, Madelain et al (submitted) developed a mathematical model to fit the longitudinal SIV RNA data in this cohort of animals. The best fit to the data was obtained by using a model including an immune-response-mediated infected-cell elimination compartment where the loss rate of productively infected cells increased over time. Interestingly, the pattern of increase in cell loss rates (based on the analysis of SIV RNA only) nicely matched in most animals the changes in the capacity of CD8^+^ T-cells to suppress infection that were obtained experimentally. Moreover, a *post hoc* positive correlation was found between the theoretical immune-response-mediated infected-cell elimination rate and the experimental CD8^+^ T-cell-mediated SIV-suppressive activity, but not with the frequency of SIV-specific CD8^+^ T-cells.

Therefore, our results exposed a disconnection between the development of SIV-specific CD8^+^ T-cells producing cytokines and cytolytic molecules and the ability of these cells to suppress SIV infection, as measured *ex vivo* (Figure 5E, Figure S6). SIV-specific CD8^+^ T-cell frequencies increased sharply as the initial viremia began to fall and remained high for the duration of the study in CyMs irrespectively of their level of viremia. However, the substantial decline in viremia to levels below 400 copies/mL in SICs coincided with the raise of SIV-suppressive activity *ex vivo*. The increase of CD8^+^ T-cell-mediated SIV-suppressive activity was delayed in the late controller #BL669 and #BO413, but nonetheless preceded optimal control of viremia in these CyMs. A very strong capacity of CD8^+^ T-cells to suppress SIV was observed at day 36 in the LN from two animals (BA209 and BC657) that did not show such capacity in the blood (Figure 5A). Only one animal (29925) did not develop any detectable SIV suppressive activity during our follow up. This animal had the weakest peak of viremia (1 log lower than any other) and achieved the fastest control of viremia. Whether a very rapid or local development of the CD8^+^ T-cell suppressive capacity may have occurred or other mechanisms were associated with control of viremia in this animal remains unknown (Figure S6). At the time of euthanasia, superior CD8^+^ T-cell-mediated SIV-suppressive activity was detected in a vast majority of SICs across all anatomical compartments, with the exception of bone marrow (Figure 5A), which nonetheless harbored SIV-specific CD8^+^ T-cells at frequencies comparable to other tissues (Figure 4A). Thus, although abundant, SIV-specific CD8^+^ T-cells induced during primary SIV infection had limited SIV suppressive capacity when compared to cells found at later time points in SICs (Figure 5E).

To confirm that the capacity of CD8^+^ T-cells to suppress ex vivo SIV infection did not increase in VIRs, we analyzed this activity in an additional group of 14 non-M6 CyMs infected intravenously (*i.v.*) with 1,000AID_50_ of SIVmac251 and characterized by high setpoint viremia (ANRS pVISCONTI study). In these animals, the CD8^+^ T-cell-mediated SIV-suppressive activity also remained modest throughout the follow-up (Figure S8A). The combined analysis of the CD8+ T-cells from all VIR CyMs (n=4 50AID_50_ + n=14 1,000AID_50_) exposed early significant differences in the CD8^+^ T-cell-mediated SIV-suppressive activity when compared to the SICs (Figure S8B). Moreover, early initiation (day 28 post-infection) of antiretroviral treatment in another group of CyM inoculated with 1,000AID_50_ of SIVmac251 sharply decreased viremia and CD8^+^ T-cell activation levels (Figure S8C) but did not change the capacity of CD8^+^ T-cells from these animals to suppress infection *ex vivo*, which remained extremely weak (Figure S8C). These results, which are in agreement with our previous observations in early treated HIV-infected individuals (Lecuroux et al., 2013), show that low SIV suppressive capacity during acute infection was neither a consequence of strong activation of these cells *in vivo* nor of high antigen burden.

Collectively, our results show that the capacity of SIV-specific CD8^+^ T-cells to suppress infection *ex vivo* was a genuine quality that progressively amplifies in SICs. Our results further uncover a temporal link between the acquisition by CD8^+^ T-cells of potent capacity to suppress infection and sustained control of SIV.

### Acquisition of CD8^+^ T-cell-mediated SIV-suppressive activity in SICs occurs independently of MHC haplotype

Our primary intention in this study was to explore the mechanisms underlying natural control of SIV infection, independently of MHC background or infectious dose. However, as expected (Bruel et al., 2015), inoculation with low-dose virus and carriage of the protective M6 haplotype independently favored spontaneous control of viremia below 400 copies/mL in CyMs in our study (Supplemental Table 3). We therefore evaluated whether these parameters influenced the dynamics of control and the development of the CD8^+^ T-cell response upon infection. We found that CD4^+^ T-cells and the levels of cell-associated SIV DNA similarly evolved in the blood and PLNs from M6 and non-M6 controllers (Figure S9A). There was just a tendency for M6 controllers vs non-M6 controllers to better recovery of CD4^+^ T-cells in blood at the end of the study (p=0.07). Similarly, we did not find important differences between M6 and non-M6 controllers in their development of SIV-specific CD8^+^ T-cell responses (Figure S9B). M6 and non-M6 controllers developed similar frequencies of SIV-responding cells during acute infection that were maintained during the follow up. Of note, the capacity of CD8^+^ T-cells to suppress *ex vivo* SIV infection of CD4^+^ T-cells progressively increased in both M6 and non-M6 controllers. The only difference that we could appreciate was a faster acquisition (day 36 *p.i.*) of CD8^+^ T-cell mediated SIV suppressive activity in the PLN from M6 SICs versus non-M6 SICs (Figure S9B). Intriguingly, non-M6 SICs had higher frequencies of SIV responding CD8^+^ T-cells in this tissue at the same time point. Overall these results show that while the M6 background gave a selective advantage to CyMs to control infection in conditions of higher viral inoculum, this MHC haplotype was not indispensable for the acquisition of SIV suppressive capacity by CD8^+^ T-cells, which occurred both in M6 and non M6 SICs. The results are in agreement with the observations in HIV controllers. Although cohorts of HICs are enriched in individuals carrying protective HLA class I alleles (mainly HLA-B*57, B*27), many HICs do not carry protective HLA class I alleles but have CD8^+^ T-cells with strong HIV suppressive capacity *ex vivo* (Lecuroux et al., 2014). Therefore, the development of efficient CD8^+^ T-cell responses with antiviral activity is a characteristic of most HICs/SICs, independently of their MHC background.

### Skewed maturation of central memory SIV-specific CD8^+^ T-cells is associated with defective acquisition of SIV-suppressive activity

To dissect the phenotypic correlates of *ex vivo* measured antiviral potency, we analyzed the differentiation status of SIV-specific CD8^+^ T-cells using selected markers in conjunction with MHC class I tetramers (Figure S10, S11). Tetramer-binding SIV-specific CD8^+^ T-cells were detected in all CyMs, displayed early similar differentiation profiles in SICs and VIRs, but evolved differently, such that higher frequencies of central memory (CM) SIV-specific CD8^+^ T-cells were present in SICs versus VIRs on day 105 *p.i.* (p = 0.018) and day 535 *p.i.* (p = 0.013) (Figure 6A, B).

**Figure 6.**
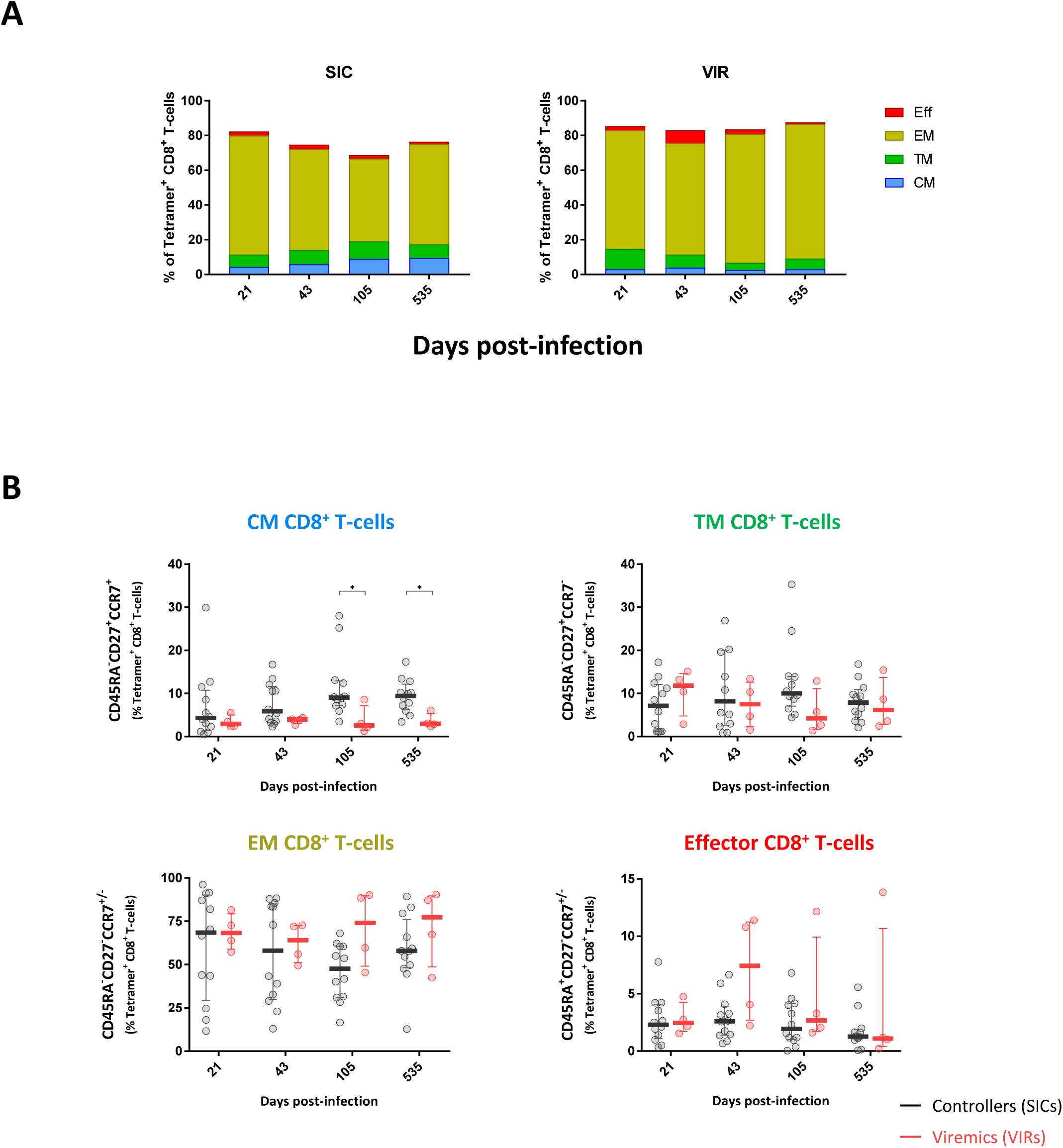
SIV controllers maintain higher frequencies of SIV-specific central memory CD8^+^ T-cells during chronic infection than viremic CyMs. (**A**) Doughnut charts showing median percent frequencies of SIV-specific CD8^+^ T-cells in each phenotypically-defined subset in SIV controllers (upper panels) and viremic CyMs (lower panels). Light blue, central memory (CM); green, transitional memory (TM); yellow, effector memory (EM); red, effector (Eff). (**B**) Evolution of CM, TM, EM, and Eff SIV-specific CD8^+^ T-cells in SIV controllers (black) and viremic CyMs (red). Results are shown as percent frequencies of tetramer-binding CD8^+^ T-cells. *p < 0.05; Mann-Whitney U-test.

In further analyses, we found that higher frequencies of SIV-specific CD8^+^ T-cells from SICs expressed the IL-7 receptor CD127, which is associated with cell survival and memory responses (Schluns et al., 2000), whereas higher frequencies of SIV-specific CD8^+^ T-cells from VIRs expressed the transcription factor T-bet, which is associated with cellular differentiation and effector functionality (Sullivan et al., 2003; Szabo et al., 2002) (Figure 7A). These differences appeared since primary infection and became statistically significant at later time points (Figure 7A). Expression levels of CD127 and T-bet also varied as a function of differentiation among SIV-specific CD8^+^ T-cells from SICs and VIRs (Figure 7B). In particular, CM and transitional memory (TM) SIV-specific CD8^+^ T-cells expressed lower levels of T-bet throughout the course of infection in SICs *versus* VIRs, whereas CM SIV-specific CD8^+^ T-cells tended to express higher levels of CD127 during chronic infection in SICs *versus* VIRs.

**Figure 7.**
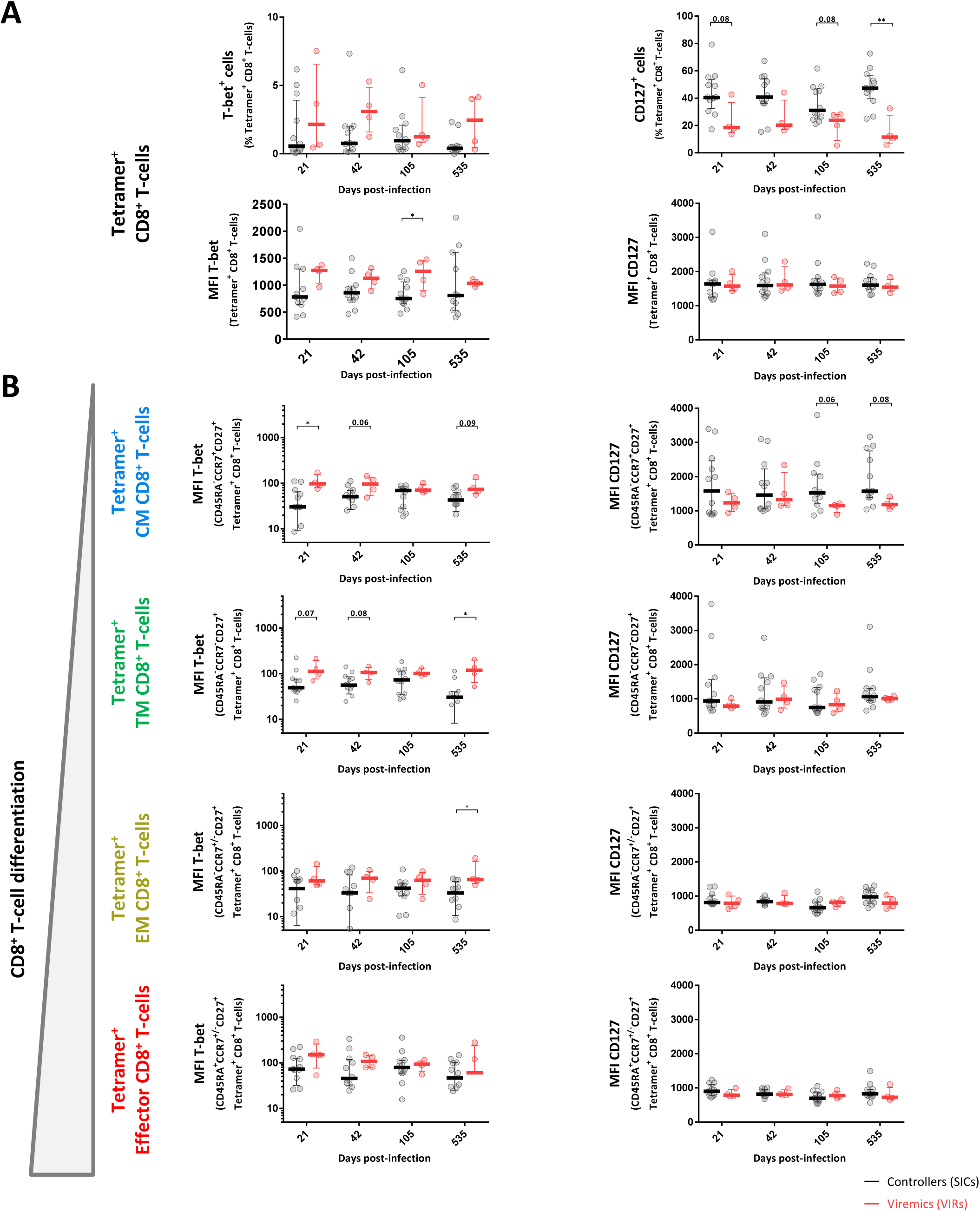
Altered maturation of central memory SIV-specific CD8^+^ T-cells in viremic CyMs. (**A**) Dynamics of T-bet (left panels) and CD127 expression (right panels) among SIV-specific CD8^+^ T-cells in SIV controllers (black) and viremic CyMs (red). (**B**) Dynamics of T-bet (left) and CD127 expression (right) among central memory (CM), transitional memory (TM), effector memory (EM), and effector (Eff) SIV-specific CD8^+^ T-cells in SIV controllers (black) and viremic CyMs (red). *p < 0.05, **p < 0.01; Mann-Whitney U-test.

Accordingly, negative correlations were observed during primary infection and at euthanasia between the expression levels of CD127 on SIV-specific CD8^+^ T cells and plasma viral loads (Figure 8A). Of note, the levels of CD127 correlated positively with CD8^+^ T-cell-mediated SIV-suppressive activity at the same time points (Figure 8B). On the contrary, negative correlations were observed during primary infection and at euthanasia between CD8^+^ T-cell-mediated SIV-suppressive activity and the contemporaneous frequencies of T-bet^+^CD127^−^ SIV-specific CD8^+^ T-cells (Figure 8C) and between CD8^+^ T-cell-mediated SIV-suppressive activity and expression levels of T-bet among CM SIV-specific CD8^+^ T-cells (Figure 8D).

**Figure 8.**
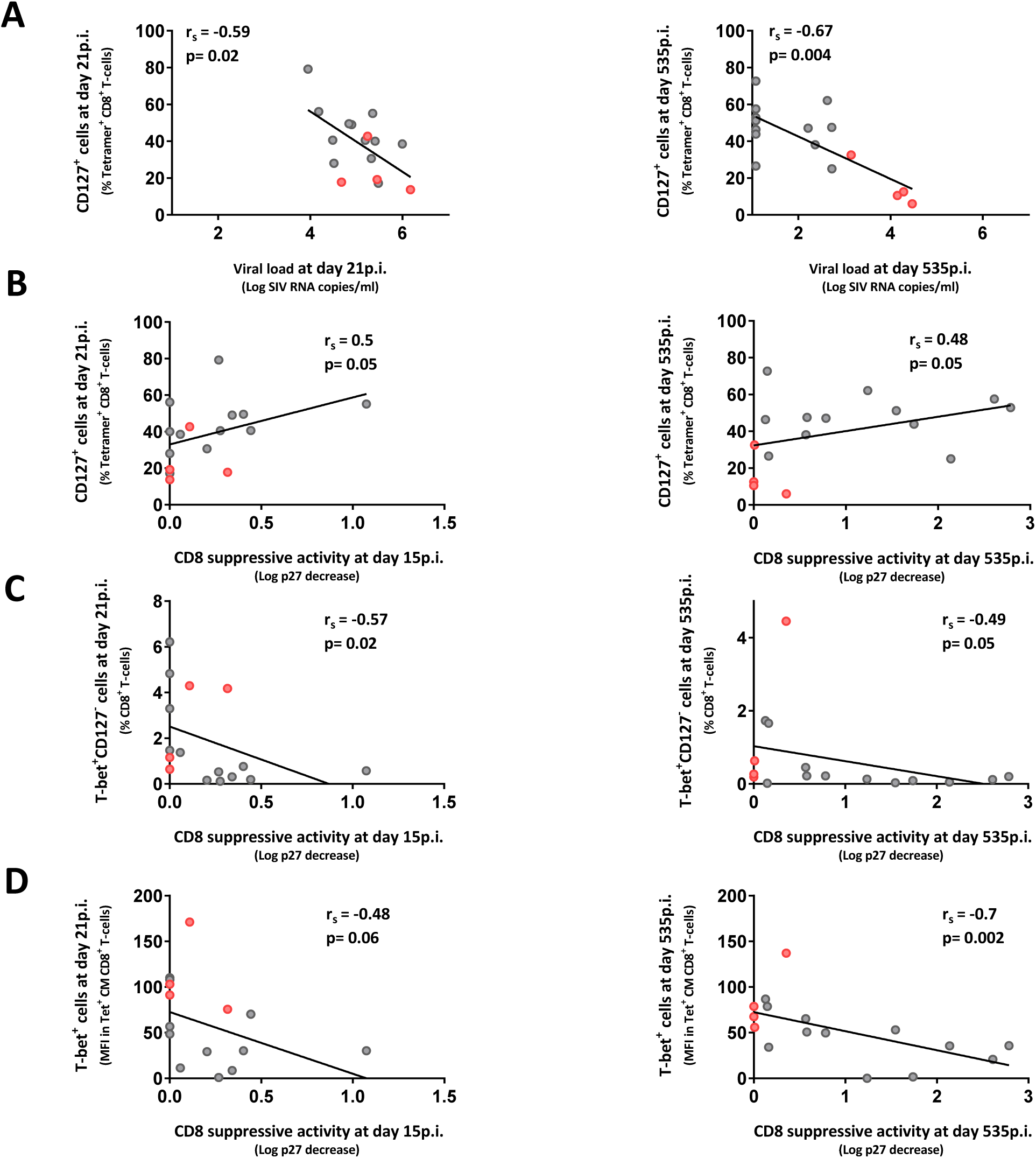
Skewed maturation of central memory SIV-specific CD8^+^ T-cells is associated with defective acquisition of SIV-suppressive activity. Spearman correlations between CD127+ SIV specific CD8^+^ T cell frequencies and viral loads (**A**) and CD8^+^ T-cell-mediated SIV-suppressive activity (**B**) during acute (left panel) and chronic infection (right panel). Spearman correlations between T-bet^+^ CD127^−^ SIV-specific CD8^+^ T-cell frequencies (**C**) or T-bet expression levels in central memory SIV-specific CD8^+^ T-cells (**D**) and CD8^+^ T-cell-mediated SIV-suppressive activity during acute (left panel) and chronic infection (right panel). Grey symbols, SIV controllers; red symbols, viremic CyMs.

Collectively, these results suggest that SIV-specific CM CD8^+^ T-cells are primed for survival in SICs, enabling long-term memory, sustained antiviral activity and viral control, whereas the corresponding SIV-specific CD8^+^ T-cells in VIRs adopt a skewed phenotype associated with cellular differentiation and suboptimal antiviral activity.

## DISCUSSION

The data presented in this study provide new insights into the immune correlates of natural control of SIV. Although SIV-specific CD8^+^ T-cells were generated during acute infection with equivalent dynamics and global frequencies in all CyMs, preventing discrimination between SICs and VIRs, antiviral efficacy *ex vivo* developed progressively over time and was associated with spontaneous SIV control. This dichotomy was underpinned by distinct early memory programs within the SIV-specific CD8^+^ T-cell pool. Collectively, these findings identify a cohesive set of immunological parameters that associate with effective and sustained control of SIV.

To monitor the establishment of natural control prospectively, we took advantage of previous reports showing that carriage of the MHC haplotype M6 and *i.r.* inoculation with low-dose (5AID_50_) virus independently favor spontaneous control of SIVmac251 infection in CyMs (Aarnink et al., 2011; Bruel et al., 2015; Mee et al., 2009). Our results corroborate previous reports. In particular, although the presence M6 haplotype favored more frequent and more rapid control of infection among animals receiving a high dose of the virus (50AID_50_) (Supplemental Table 3), no significant differences were observed in the dynamics of SIV control in M6 and non-M6 controllers. At the time of euthanasia, a higher proportion of CD4^+^ T-cells and lower cell-associated SIV-DNA levels were found in multiple tissues from SICs *versus* VIRs, demonstrating systemic control of SIV. These differences were much more subtle during primary infection. However, PLNs from SICs harbored approximately 10-fold less SIV-DNA in the acute phase than PLNs from VIRs. In addition, the frequency of CD4^+^ T-cells were maintained close to baseline throughout the course of the study in PLNs, but not in blood or RBs, from SICs. These observations suggest that early containment of viral replication in lymph nodes (Buggert et al., 2018; Reuter et al., 2017) may be a key event for subsequent immune control of SIV.

In line with previous studies in humans (Lecuroux et al., 2013; Ndhlovu et al., 2015; Trautmann et al., 2012) and non-human primates we observed early and robust expansions of SIV-specific CD8^+^ T-cells in all CyMs. However, the functional profiles and overall frequencies of SIV-specific CD8^+^ T-cells (as determined by intra cellular cytokine staining upon SIV antigen stimulation) during the acute phase of infection were largely equivalent in SICs and VIRs, and neither parameter correlated with subsequent determinations of plasma VL. Similarly, the functional profiles and overall frequencies of SIV-specific CD8^+^ T-cells during the chronic phase of infection were largely equivalent in SICs and VIRs, although polyfunctionality (defined as the capacity to produce simultaneously several cytokine and/or degranulate) was impaired at the time of euthanasia in VIRs. These results suggest that differences in polyfunctionality found during chronic infection are a surrogate marker of viral replication rather than an accurate determinant of antiviral efficacy, although low number of animals in the VIR group may limit statistical power.

The capacity of CD8^+^ T-cells to suppress infection of autologous CD4^+^ T-cells directly *ex vivo* is a particular feature of HICs (Almeida et al., 2009; Angin et al., 2016; Buckheit et al., 2012; Julg et al., 2010; Saez-Cirion et al., 2007; Saez-Cirion et al., 2009; Tansiri et al., 2015) that is mediated by the rapid elimination of infected CD4+ T-cells (Saez-Cirion et al., 2007). Irrespective of subsequent outcome, we detected relatively weak CD8^+^ T-cell-mediated SIV-suppressive activity during primary infection, despite the vigorous mobilization of SIV-specific CD8^+^ T-cells. This observation parallels our previous findings in the setting of HIV (Lecuroux et al., 2013) and point to limited antiviral potential of CD8^+^ T-cell responses generated during primary infection. However, a remarkable negative correlation was already observed between the CD8^+^ T-cell-mediated SIV-suppressive activity and viremia at this early time point, showing early temporal association of this antiviral activity and reduction of viremia. Of note, this SIV-suppressive capacity of CD8^+^ T-cells increased progressively over a period of weeks in some animals, carrying or not the protective MHC haplotype M6, and correlated temporally with the establishment of viral control. At the time of euthanasia, these highly potent antiviral CD8^+^ T-cells were present in all tissues, with the exception of bone marrow. It is important to notice that CD8^+^ T-cell-mediated SIV suppression was very weak also in LN during the first weeks following infection but increased over time in SICs. Therefore, the increase in the capacity of CD8^+^ T-cells to suppress infection that we observed in this study was not the result of the recirculation of CD8^+^ T-cells from lymph nodes once control was established but a genuine progressive augmentation of the antiviral potential of the cells. The development of potent antiviral CD8^+^ T-cells is therefore a *bone fide* correlate of sustained control of SIV.

The divergent antiviral properties of SIV-specific CD8^+^ T-cells in SICs *versus* VIRs were associated with early differences in the expression of CD127 and T-bet, especially within the less differentiated memory pools (CM and TM). In particular, higher frequencies of SIV-specific CM CD8^+^ T-cells expressed CD127 in SICs, whereas higher frequencies of SIV-specific CM and TM CD8^+^ T-cells expressed T-bet in VIRs. These differences became more pronounced throughout the course of infection. Studies in mice have shown that decreased expression of T-bet among memory CD8^+^ T-cells allows the establishment of long-lived CD127^hi^ cells, which maintain the capacity to proliferate and control successive infections (Joshi et al., 2007; Joshi et al., 2011). Accordingly, our data suggest that SICs develop true memory-like SIV-specific CD8^+^ T-cell responses, which is key for the acquisition of antiviral ability, whereas VIRs develop SIV-specific memory CD8^+^ T-cell responses skewed towards more effector-like characteristics. In line with this supposition, the proportion of CD127^+^ SIV-specific CD8^+^ T cells during acute infection (day 15 *p.i.*) and at euthanasia correlated positively with CD8^+^ T-cell-mediated SIV-suppressive activity at the corresponding time points, while the frequencies of T-bet^+^ CD127^−^ SIV-specific CD8^+^ T-cells and the expression levels of T-bet among CM SIV-specific CD8^+^ T-cells during acute infection correlated negatively with CD8^+^ T-cell-mediated SIV-suppressive activity. These findings are broadly consistent with several previous reports describing immune profiles that associate with the control of viremia in HICs during chronic infection. Favorable characteristics include high frequencies of CD57^+^ eomesodermin^hi^ HIV-specific CD8^+^ T-cells with superior proliferative capacity, increased expression levels of CD127, and intermediate expression levels of T-bet (Simonetta et al., 2014), and high frequencies of HIV-specific CD8^+^ T-cells with the capacity to upregulate T-bet, granzyme B, and perforin in response to antigen encounter (Hersperger et al., 2011b; Migueles et al., 2008).

In a recent single cell study (Angin et al., 2019b), we also found differences in the program of HIV-specific CM CD8^+^ T-cells from HIV controllers and non-controllers on cART: whereas HIV-specific CM CD8^+^ T-cells from HIC upregulated the expression of effectors genes linked with mTORC2 activation and cell survival (including CD127), central memory cells from non-controllers had a skewed profile associated with mTORC1 activation (including T-bet) and glycolysis. This was traduced in a dependency on glucose of HIV-specific CD8+ T-cells from non-controllers to react to HIV antigens, while HIV-specific CD8+ T-cells from controllers were characterized by metabolic plasticity and being able to exert their function even in conditions of glucose deprivation. Of note, these differences in the metabolic program of cells from controllers and non-controllers could also be recapitulated with SIV-specific CD8+ T-cells from SICs and VIR CyMs from the present study (Angin et al., 2019b), further corroborating the validity of our CyM model to study the development of the protective CD8+ T-cell responses characteristics of HIV/SIV controllers. The present results extend these observations and support a key role for long-lived memory responses in the control of SIV. Importantly, our data also show that distinct memory responses are formed early after infection, potentially reflecting different priming conditions. Interestingly, although the antiviral activity of CD8^+^ T-cells increased over time in SICs, we already found a negative correlation between this activity and the plasma viremia at day 15. On this basis, we propose that the amplification of potent antiviral activity maters in long term control and is the result of a maturation process, the trajectory of which is linked to early optimal programming of the CD8^+^ T-cell memory compartment.

It remains unclear which factors are required to encourage the development of memory CD8^+^ T-cell responses that provide optimal protection against HIV/SIV. In some viral infections, expression of T-bet is tightly regulated by cytokines, such as IL-12 (Rao et al., 2012; Takemoto et al., 2006). Low levels of inflammation may therefore favor the emergence of long-lived memory CD8^+^ T-cells. It is also interesting to note that maturation through persistent or repeated exposure to antigen can drive the selection of specific clonotypes bearing high-affinity T-cell receptors (TCRs) (Busch and Pamer, 1999; Ozga et al., 2016; Price et al., 2005) which have been shown to suppress HIV replication more efficiently than clonotypes targeting the same antigen via low-affinity TCRs (Almeida et al., 2007; Almeida et al., 2009; Ladell et al., 2013). Increase in antigen sensitivity over time would be compatible with the progressive increase in antiviral potency that we observed for the CD8^+^ T-cells from controllers in our study.

A recent study in the LCMV murine model of infection has shown that memory CD8^+^ T-cell responses expressing the transcription factor TCF1 developed during chronic infection (in an immunosuppressive environment) have a distinct molecular program, resist contraction, had increased long-term functionality, are less prone to exhaustion and are thus critical for controlling ongoing viral replication; in contrast, memory cells that are developed at the onset of infection (in a pro-inflammatory environment) become short-term effectors and are rapidly exhausted (Snell et al., 2018). Accordingly, we suggest that balanced inflammatory responses (Barouch et al., 2016) arising as a consequence of lower viral burdens in lymph nodes during acute infection in SICs might facilitate antigen-specific priming events associated with optimal memory programs (Ozga et al., 2016) and minimize the loss of CD4^+^ T-cells, which provide helper functions that are critical for the development of long-lived memory CD8^+^ T-cells (Khanolkar et al., 2004).

Collectively, the data presented here underscore the importance of early host-pathogen interactions in the development of adaptive immunity and reveal an optimal maturation pathway associated with the generation and maintenance of potent and sustained antiviral CD8^+^ T-cell responses, which in turn dictate the outcome of infection with SIV.

## METHODS

### Ethical statement

Cynomolgus macaques (CyMs, *Macaca fascicularis*) were imported from Mauritius and housed in facilities at the *Commissariat à l’Energie Atomique et aux Energies Alternatives* (CEA, Fontenay-aux-Roses, France). All non-human primate studies at the CEA are conducted in accordance with French National Regulations under the supervision of National Veterinary Inspectors (CEA Permit Number A 92-03-02). The CEA complies with the Standards for Human Care and Use of Laboratory Animals of the Office for Laboratory Animal Welfare under Assurance Number #A5826-01. All experimental procedures were conducted according to European Directive 2010/63 (Recommendation Number 9). The SIC and pVISCONTI studies were approved and accredited under statement number A13-005 and A15-035 by the ethics committee “*Comité d’Ethique en Expérimentation Animale du CEA”*, registered and authorized under Number 44 and Number 2453-2015102713323361v2 by the French Ministry of Education and Research. CyMs were studied with veterinary guidance, housed in adjoining individual cages allowing social interactions, and maintained under controlled conditions with respect to humidity, temperature, and light (12 hour light/12 hour dark cycles). Water was available *ad libitum*. Animals were monitored and fed once or twice daily commercial monkey chow and fruit by trained personnel. Environmental enrichment was provided including toys, novel foodstuffs, and music under the supervision of the CEA Animal Welfare Body. Experimental procedures (animal handling, viral inoculations, and samplings) were conducted after sedation with ketamine chorhydrate (Rhone-Merieux, Lyon, France, 10 mg/kg). Tissues were collected at necropsy: animals were sedated with ketamine chlorhydrate 10 mg/kg) then humanely euthanized by intravenous injection of 180 mg/kg sodium pentobarbital.

### Animals and SIV infection

A total of 16 healthy adult male CyMs (median age = 6.8 years at inclusion, IQR = 5.8–7.2) were selected for this study on the basis of MHC haplotype (M6^+^, n = 6; M6^−^, n = 10) *(34)*. CyMs were inoculated *i.r.* with either 5AID_50_ or 50AID_50_ of uncloned SIVmac251 (A.M. Aubertin, Université Louis Pasteur, Strasbourg, France). The following experimental groups were studied: (i) M6^−^ CyMs inoculated *i.r.* with 5AID_50_ (non-M6 5AID_50_, n = 4); (ii) M6^+^ CyMs inoculated *i.r.* with 50AID_50_ (M6 50AID_50_, n = 6); and (iii) M6^−^ CyMs inoculated *i.r.* with 50AID_50_ (non-M6 50AID_50_, n = 6). Animals were monitored for 18 months post-infection.

The outcome of infection generally matched expectations based on previous studies for each experimental group (Figure S12, Supplemental Table 1). Only one M6^+^ CyM (31041) was unable to control viremia below 400 copies/mL. This animal was homozygous for MHC class I (Supplemental Table 1), which intrinsically limits immune control of HIV/SIV (Carrington et al., 1999; O’Connor et al., 2010). The dynamics of viral replication during acute infection were very similar in the three experimental groups, with peak VLs of 5.9, 6.4, and 6.3 log SIV-RNA copies/mL of plasma on day 14 *p.i.* for non-M6 5AID_50_, M6 50AID_50_, and non-M6 50AID_50_ CyMs, respectively (Supplemental Table 1).

CyMs in the pVISCONTI study (median age = 5 years at inclusion, IQR = 4.1–5.3) were inoculated with 1000 AID_50_ of uncloned SIVmac251 through the intravenous route. None of these animals carried the M6 haplotype. An antiretroviral regimen containing emtricitabine (FTC), dolutegravir (DTG), and the tenofovir prodrug tenofovir-disoproxil-fumarate (TDF), co-formulated as a once daily subcutaneous injection, was initiated at day 28 post-inoculation in 6 animals. TDF was administered at 5.1 mg/kg, FTC at 40 mg/kg and DTG at 2.5 mg/kg.

### Blood collection and processing

Peripheral blood was collected by venous puncture into Vacutainer Plus Plastic K3EDTA Tubes or Vacutainer CPT Mononuclear Cell Preparation Tubes with Sodium Heparin (BD Biosciences). Complete blood counts were monitored at all time points from the Vacutainer Plus Plastic K3EDTA Tubes. Plasma was isolated from Vacutainer Plus Plastic K3EDTA Tubes by centrifugation for 10 min at 1,500 g and stored at –80 °C. Peripheral blood mononuclear cells (PBMCs) were isolated from Vacutainer CPT Mononuclear Cell Preparation Tubes with Sodium Heparin according to manufacturer’s instructions (BD Biosciences), and red blood cells were lysed in ACK (NH_4_Cl 0.15 M, KHCO_3_ 10 mM, EDTA 0.1 mM, pH 7.4).

### Tissue collection and processing

Axillary or inguinal lymph nodes (PLNs), rectal biopsies (RBs) and broncho-alveolar lavages (BAL) were collected longitudinally from each animal at the indicated time points. In addition, bone marrow, spleen, mesenteric lymph nodes (MLNs), duodenum, jejunum, ileum and colon were collected at necropsy. Tissue samples were snap-frozen in liquid nitrogen for storage at –80 °C or collected in RPMI medium at 2–8 °C. At each time point a complete PLN group was collected. LNs were washed and cells were freshly isolated in RPMI medium upon mechanical disruption with a GentleMACS dissociator as recommended by the manufacturer (Miltenyi Biotec). Cell suspension was filtered (70µm), then red blood cells were lysed in ACK. RB lymphocytes were obtained from approximately 4 mm^2^ of rectal mucosa. Colonic lymphocytes were obtained from mucosa taken from approximately 10 cm of tissue. RBs and colonic tissue were washed extensively in R10 medium (RPMI medium supplemented with 10% fetal calf serum and penicillin/neomycin/streptomycin), then digested for 45 minutes with collagenase II prior to mechanical disruption. Lymphocytes were isolated over a Percoll 67/44 gradient (Sigma-Aldrich). Bone marrow cells were purified using Lymphocyte Separation Medium (Lonza Bioscience) diluted to 90% in DPBS, centrifuged for 20 minutes at 350 g, and separated from red cells in ACK. Spleen cells were processed by mechanical disruption in RPMI medium using a GentleMACS™ Dissociator (Miltenyi Biotec), purified as described for bone marrow cells, and separated from red cells in ACK. Total cells were immediately designated to T-cell activation and proliferation analyses by flow cytometry, CD4^+^ and CD8^+^ T-cells separation with magnetic beads for antiviral activity assay and the remaining cells were frozen for further assessment of cytokine production by ICS or tetramer analyses.

### Quantification of plasma viral load

Plasma viremia was monitored longitudinally in all animals using quantitative real-time RTqPCR with a limit of detection of 12.3 copies/mL (Angin et al., 2019a). Viral RNA was prepared from 100 μl of cell-free plasma. Quantitative RT-PCR was performed with a SuperScript III Platinum One-Step qRT-PCR Kit (Thermofisher) in a CFX96 Touch Real-Time PCR Detection System (BioRad) under the following conditions: 12.5 μl of 2X reaction mixture, 0.5 μl of RNaseOUT (40U/μl), 0.5 μl of Superscript III reverse transcriptase/Platinum Taq DNA Polymerase, 1 μl of each primer (125 μM), 0.5 μl of the fluorogenic probe (135 μM), and 10-μl of RNA elution samples. The probe and primers were designed to amplify a region of SIVmac251 *gag*. Forward primer was 5’-GCAGAGGAGGAAATTACCCAGTAC-3’ (24 bp) and reverse primer was 5’-CAATTTTACCCAGGCATTTAATGTT-3’ (25 bp). The TaqMan probe sequence was 5’-FAM-TGTCCACCTGCCATTAAGCCCGA-BHQ1-3’ (23 bp). This probe had a fluorescent reporter dye, FAM (6-carboxyfluorescein), attached to its 5’ end and the quencher BHQ1 (Black Hole Quencher 1) attached to its 3’ end. Samples were heated for 30 min at 56°C and 5 min at 95°C, followed by 50 cycles of 95°C for 15 s and 60°C for 1 min.

### Quantification of SIV-DNA

Total DNA was extracted from purified CD14^+^ alveolar macrophages, buffy coats and snap-frozen tissues. CD14^+^ alveolar macrophages were purified by positive selection using antibody-coated magnetic beads following manufacturer’s instructions (Miltenyi Biotec). Purity was checked by flow cytometry (Figure S1A, upper panel). Snap-frozen tissues were mechanically disrupted with a MagNA Lyser (Roche Diagnostics). DNA extraction was performed using a QIAamp DNA Blood Mini Kit (Qiagen) following manufacturer’s instructions. SIV-DNA was quantified using an ultrasensitive quantitative real-time PCR. For blood samples, 150,000 cells were analyzed for each SIV-DNA PCR. Due to sample size limitations, for rectal biopsies and bronchoalveolar lavages 50,000 and 20,000 cells per PCR were tested, respectively. All amplifications were performed on 2–4 replicates. The cell line SIV1C, which contains 1 copy of SIV integrated/cell, was used as a standard for quantification. 1 μg of DNA was considered to be equivalent to 150,000 cells. Amplification was performed using primers and a probe located in the *gag* region. The *CCR5* gene was used to normalize results per million cells. Results were then adjusted by the frequencies of CD4^+^ T-cells in blood and tissues, when available. The limit of quantification was 2 copies/PCR. Primers and probes were: SIV *gag* F: 5’-GCAGAGGAGGAAATTACCCAGTAC-3’; SIV *gag* R: 5’-CAATTTTACCCAGGCATTTAATGTT-3’; SIV *gag* probe: 5’-FAM-TGTCCACCTGCCATTAAGCCCGA-BHQ1-3’; *CCR5* F: an equimolar mix of 5’-CAACATGCTGGTCgATCCTCAT-3’ and 5’-CAACATACTGGTCGTCCTCATCC-3’; *CCR5* R: 5’-CAGCATAGTGAGCCCAGAAG-3’; and *CCR5* probe: 5’-HEX-CTGACATCTACCTGCTCAACCTG-BHQ1-3’.

### Measurement of T-cell activation and proliferation

T-cell activation and proliferation were assessed using fresh PBMCs and tissue cell suspensions. Blood samples were treated with FACS Lysing Solution (BD Biosciences). Cells were surface stained for CD3, CD4, CD8, CD38, CD45, CCR5, and HLA-DR, fixed/permeabilized using a Cytofix/CytoPerm Kit (BD Biosciences), and stained intracellularly for Ki-67. The following antibodies used were: anti-CD3–PE (clone SP34-2, BD Biosciences), anti-CD4–PerCP-Cy5.5 (clone L200, BD Biosciences), anti-CD8–BV650 (clone RPA-T8, BioLegend), anti-CD38–FITC (clone AT-1, StemCell Technologies), anti-CD45–V500 (clone D058-1283, BD Biosciences), anti-CCR5–APC (clone 3A9, BD Biosciences), anti-HLA-DR–APC-H7 (clone G46-6, BD Biosciences), and anti-Ki-67–AF700 (clone B56, BD Biosciences). Data were acquired using an LSRII flow cytometer (BD Biosciences) and analyzed with FlowJo software version 10 (TreeStar Inc.).

### Intracellular cytokine staining

Frozen PBMCs, PLN cells, bone marrow cells, splenocytes and MLN cells were thawed, resuspended at 1 × 10^6^/mL in R20 medium, and stored overnight at 37 °C. Cells were then stimulated with a pool of 24 optimal SIV peptides (8-10 amino acids long) (2 μg/mL each, Supplemental Table 2) or with a pool of 125 overlapping SIV Gag 15-mer peptides (2 μg/mL each, NIH AIDS Reagent Program, SIVmac239 Gag Peptide Set #12364) in the presence of anti-CD28 (1 μg/mL, clone L293, BD Biosciences) and anti-CD49d (1 μg/mL, clone 9F10, BD Biosciences) and stained with anti-CD107a (clone H4A3, BD Biosciences) for 30 minutes prior to the addition of GolgiStop (1 μL/mL, BD Biosciences) and brefeldin A (BFA, 5 μg/mL, Sigma-Aldrich). Costimulatory antibodies alone were used as a negative control, and concanavalin A (5 µg/mL, Sigma-Aldrich) was used as a positive control. Cells were incubated for a total of 6 hours. After washing, cells were surface stained for CD3, CD4, and CD8, fixed/permeabilized using a Cytofix/CytoPerm Kit (BD Biosciences), and stained intracellularly for IFNγ, TNFα, and IL-2. The following antibodies were used: anti-CD107a–V450 (clone H4A3, BD Biosciences), anti-CD3–AF700 (clone SP34-2, BD Biosciences), anti-CD4–PerCP-Cy5.5 (clone L200, BD Biosciences), anti-CD8–APC-Cy7 (clone RPA-T8, BD Biosciences), anti-IFNγ–PE-Cy7 (clone B27, BD Biosciences), anti-IL-2–PE (clone MQ1-17H12, BD Biosciences), and anti-TNFα–PE-CF594 (clone Mab11, BD Biosciences). Data were acquired using an LSRII flow cytometer (BD Biosciences) and analyzed with FlowJo software version 10 (TreeStar Inc.). Results were corrected for background by subtracting the peptide stimulated response from the negative (no peptide) control. Negative responses were given an arbitrary value of 0.001. All data are represented. A representative flow cytometry gating strategy used to analyze cytokine production via intracellular staining after peptide stimulation is shown in Figure S13.

### MHC class I tetramer staining

Biotinylated complexes of Nef RM9 (RPKVPLRTM)–Mafa A1*063:02, Gag GW9 (GPRKPIKCW)–Mafa A1*063:02, and Vpx GR9 (GEAFEWLNR)–Mafa B*095:01 were produced as described previously*(53)*. The corresponding tetramers were generated via stepwise addition of APC-conjugated streptavidin (Thermo Fisher Scientific). Frozen PBMCs were stained with the pool of these tetramers for 30 minutes at 37 °C, washed, and surface stained for CD3, CD4, CD8, CD14, CD20, CD27, CD45RA, CCR7, HLA-DR, and CD127. Cells were then fixed/permeabilized using a Cytofix/CytoPerm Kit (BD Biosciences) and stained for T-bet. The following antibodies were used: anti-CD3–AF700 (clone SP34-2, BD Biosciences), anti-CD4–PerCP-Cy5.5 (clone L200, BD Biosciences), anti-CD8–APC-Cy7 (clone RPA-T8, BD Biosciences), anti-CD14–BV786 (clone M5E2, BD Biosciences), anti-CD20–BV786 (clone L27, BD Biosciences), anti-CD27–PE (clone M-T271, BD Biosciences), anti-CD45RA–PE-Cy7 (clone 5H9, BD Biosciences), anti-CCR7–PE-Dazzle594 (clone G043H7, BioLegend), anti-HLA-DR– Pacific Blue (clone G46-6, BD Biosciences), anti-CD127–FITC (clone MB15-18C9, Miltenyi Biotec), and anti-T-bet–BV711 (clone 4B10, BioLegend). Data were acquired using an AriaIII flow cytometer (BD Biosciences) and analyzed with FlowJo software version 10 (TreeStar Inc.). A representative flow cytometry gating strategy used to analyze T-cell differentiation and tetramer staining are shown in Figure S10, S11.

### Measurement of SIV-suppressive activity

Autologous CD4^+^ and CD8^+^ T-cells were purified from freshly isolated PBMCs or tissue cell suspensions by positive and negative selection, respectively, using antibody-coated magnetic beads with a RoboSep instrument (StemCell Technologies). Purified CD4^+^ T-cells were stimulated for 3 days with concanavalin A (5μg/mL, Sigma-Aldrich) in the presence of IL-2 (100 IU/mL, Miltenyi Biotec). Purified CD8^+^ T-cells were cultured in the absence of mitogens and cytokines (*ex vivo* CD8^+^ T-cells). Stimulated CD4^+^ T-cells (10^5^) were superinfected in U-bottom 96-well plates with SIVmac251 (MOI = 10^−3^) in the presence (1:1 effector to target cell ratio) or absence of e*x vivo* CD8^+^ T-cells (10^5^) from the same tissue via spinoculation for 1 hour (1,200 g at room temperature) followed by incubation for 1 hour at 37 °C. Cells were then washed and cultured in R10 medium containing IL-2 (100 IU/mL, Miltenyi Biotec). Culture supernatants were assayed on day 7 using an SIV p27 Antigen ELISA Kit (Zeptometrix). Antiviral activity was calculated as log10 (mean p27 ng/mL in SIV-infected CD4^+^ T-cell cultures without CD8^+^ T-cells) / (mean p27 ng/mL in SIV-infected CD4^+^ T-cell cultures + *ex vivo* CD8^+^ T-cells) (Saez-Cirion et al., 2010).

### Data visualization and statistical analyses

Data visualization was performed using Tableau version 2018.1.4 (Tableau Software). Statistical analyses were performed using GraphPad version 8.1.2 (Prism Software) and SigmaPlot version 12.5 (SYSTAT Software). Results are given as median with interquartile range. The non-parametric Mann-Whitney U-test was used to compare data sets between groups. Correlations were assessed by Spearman-rank analyses. Given the exploratory nature of the analyses, p values were not adjusted for multiple comparisons. All p values less than 0.05 were defined as significant.

## ACKNOWLEDGMENTS

This study was funded by the French National Agency of AIDS and Viral Hepatitis Research (ANRS) and by MSDAvenir. Additional support was provided by the Programme Investissements d’Avenir (PIA), managed by the ANR under reference ANR-11-INBS-0008, funding the Infectious Disease Models and Innovative Therapies (IDMIT, Fontenay-aux-Roses, France) infrastructure, and ANR-10-EQPX-02-01, funding the FlowCyTech Facility (IDMIT, Fontenay-aux-Roses, France). C.P. was supported by an ANRS Postdoctoral Fellowship Grant, A.M. was supported by MSDAvenir, and D.A.P. was supported by a Wellcome Trust Senior Investigator Award. We thank Benoit Delache, Brice Targat, Claire Torres, Christelle Cassan, Jean-Marie Robert, Julie Morin, Patricia Brochard, Sabrina Guenounou, Sebastien Langlois, and Virgile Monnet for expert technical assistance, Antonio Cosma for helpful discussion, Lev Stimmer for anatomopathology expertise, Delphine Desjardins and Isabelle Mangeot-Méderlé for helpful project management at IDMIT, and Christophe Joubert for veterinarian assistance at the animal facility at CEA. The SIV1C cell line was kindly provided by François Villinger. The SIVmac239 Gag Peptide Set was obtained through the NIH AIDS Reagent Program, Division of AIDS, NIAID, NIH. FTC, DTG and TDF were obtained from Gilead and ViiV healthcare though the “*IAS Towards an HIV Cure”* common Material Transfer Agreement for preclinical studies in HIV cure research.

## AUTHOR CONTRIBUTIONS

C.P. and A.M. designed and performed experiments, analyzed data, and interpreted results. V.Ma., J.G., and V.A.F. analyzed data and interpreted results. V.Mo., A.D., P.V., and N.S. performed experiments and analyzed data. E.G. and D.A.P. produced bespoke reagents. N.D.B. designed experiments, analyzed data and interpreted results. D.A.P., A.B., G.P., R.L.G., O.L., M.M.T., and C.R. interpreted results. B.V. and A.S.C. designed experiments, analyzed data, interpreted results, and supervised the study. C.P., B.V., and A.S.C. wrote the paper with assistance from A.M., V.Ma., V.Mo., A.D., P.V., N.S., D.A.P., N.D.B., R.L.G., O.L., M.M.T., C.R., J.G., and V.A.F.

## DECLARATION OF INTERESTS

The authors declare no competing interests.

