## Supplementary tables and figures for "Optimal maturation of the SIV-specific CD8^+^ T-cell response after primary infection is associated with natural control of SIV. ANRS SIC study"

### Supplemental Figures

#### Supplemental Information

**Figure S1. Levels of cell-associated SIV DNA in alveolar macrophages and gut.** (**A**) Representative example of the proportion of $CD14^+$ and $CD3^+$ cells in bronchoalveolar lavages and after positive selection of $CD14^+$ cells with magnetic beads (top). Kinetics of SIV-DNA levels in purified alveolar $CD14^+$ cells. Results are expressed as copies SIV-DNA/million $CD14^+$ cells (bottom). (**B**) Levels of SIV-DNA in duodenum, jejunum, ileum, colon and rectal biopsies at euthanasia. Results are expressed as copies SIV-DNA/million cells. Grey symbols, SIV controllers; red symbols, viremic CyMs. Median and interquartile range are shown. $^*p < 0.05$, $^{**}p < 0.01$, $^{***}p < 0.001$; Mann-Whitney U-test.

**Figure S2. Frequencies of blood $CD8^+$ T-cells producing IFN$\gamma$, IL-2 or mobilizing CD107a in response to optimal SIV peptides.** (**A**–**C**) Kinetics of IFN$\gamma^+$ (**A**)**,** IL-$2^+$ (**B**), and CD107$a^+$ (**C**) SIV-specific $CD8^+$ T-cells in blood over the course of infection and in bone marrow, spleen, and mesenteric lymph nodes at euthanasia in SIV controllers (grey) and viremic CyMs (red). Results are shown as percent frequencies among $CD8^+$ T-cells. (**D**) Kinetics of total SIV-specific $CD8^+$ T-cells in blood over the course of infection and in bone marrow, spleen, and mesenteric lymph nodes at euthanasia in SIV controllers (grey) and viremic CyMs (red). Median and interquartile range are shown.

**Figure S3. Frequencies of PLN $CD8^+$ T-cells producing IFN$\gamma$, IL-2 or mobilizing CD107a in response to optimal SIV peptides.** Kinetics of IFN$\gamma^+$ (**A**)**,** IL-$2^+$ (**B**), CD107$a^+$ (**C**), and total (**D**) SIV-specific $CD8^+$ T-cells in peripheral lymph nodes over the course of infection in SIV controllers (black) and viremic CyMs (red). Results are shown as percent frequencies among $CD8^+$ T-cells. Median and interquartile range are shown.

**Figure S4. Frequencies of polyfunctional SIV-specific $CD8^+$ T-cell in response to optimal SIV peptides.** Functional profiles of SIV-specific $CD8^+$ T-cells in blood and peripheral lymph nodes over the course of infection and in bone marrow, spleen, and mesenteric lymph nodes at euthanasia in SIV controllers (black) and viremic CyMs (red). Results are shown as percent frequencies of SIV-specific $CD8^+$ T-cells expressing the indicated number of simultaneous functions (among TNFα, IFNγ, IL-2 or mobilizing CD107a). Median and interquartile range are shown. *$p < 0.05$; Mann-Whitney U-test.

**Figure S5. Frequencies of blood $CD8^+$ T-cells responding to overlapping SIV Gag peptides.** **(A)** Kinetics of $TNF\alpha^+$, $IFN\gamma^+$, IL-$2^+$, $CD107a^+$ and total SIV-specific $CD8^+$ T-cell responses to Gag peptide pool in blood over the course of infection in SIV controllers (black) and viremic CyMs (red). Median and interquartile range are shown. Total SIV-specific $CD8^+$ T-cells were estimated as $CD8^+$ T-cells producing at least one of the above functions in response SIV peptides. **(B)** Functional profiles of SIV-specific $CD8^+$ T-cells after stimulation with Gag peptide pool in blood over the course of infection in SIV controllers (black) and viremic CyMs (red). Doughnut charts show median percent frequencies of SIV-specific $CD8^+$ T-cells expressing IFNγ, TNFα, IL-2, and/or CD107a. Colors indicate number of functions (blue, 1; green, 2; yellow, 3; red, 4). Results are shown as percent frequencies of SIV-specific $CD8^+$ T-cells expressing 3 or 4 functions among $CD8^+$ T-cells. Median and interquartile range are shown. *$p < 0.05$; Mann-Whitney U-test.

**Figure S6. Individual evolution of viremia, frequency of SIV-specific CD8+ T-cells and $CD8^+$ T-cell-mediated SIV-suppressive activity.** Dynamics of plasma VL (grey), $CD8^+$ T-cell-mediated SIV-suppressive activity (orange), and TNF$\alpha$ production by SIV-specific $CD8^+$ T-cells (blue) for each CyM. Macaque ID codes are colored as follows: green, M6 $50AID_{50}$ (*i.r.*); orange, non-M6 $5AID_{50}$ (*i.r.*); blue, non-M6 $50AID_{50}$ (*i.r.*). Macaque AV979 developed a tonsillar lymphoblastic lymphoma extending into mandibular lymph nodes and was euthanized on day 442 *p.i.*. Results are shown as log p27 decrease in the presence of $CD8^+$ T-cells for SIV-suppressive activity and percent frequencies among $CD8^+$ T-cells for TNF$\alpha$ production by SIV-specific $CD8^+$ T-cells.

**Figure S7. Capacity of $CD8^+$ T-cells to suppress SIV *ex vivo* correlates negatively with viral load.** Correlations between $CD8^+$ T-cell-mediated SIV-suppressive activity (upper panel) or TNFα production by SIV-specific $CD8^+$ T-cells (bottom panel) at day 70 *p.i.* and plasma VL at different times post infection. Grey symbols represent individual values. Linear regression (lines) and p for spearman analyses are indicated for each comparison.

**Figure S8. Evolution of plasma viral loads and capacity of $CD8^+$ T-cells to suppress SIV infection *ex vivo* in two groups of CyM infected with SIVmac251 at $1000AID_{50}$ *i.v.* (viremic untreated and starting cART at day 28 post-infection).** Animals were monitored in the context of the ANRS pVISCONTI study. **(A)** Plasma viral loads (left panel) and CD8+ T-cell-mediated SIV-suppressive activity (right panel) in viremic untreated animals. Values are shown at baseline (day -20) and different days post-infection. Individual values, median and interquartile range for 14 animals are shown. **(B)** CD8+ T-cell-mediated SIV-suppressive activity in blood over the course of infection in SIV controllers (black) and in pooled viremic

CyMs (red) infected with SIVmac251 at $50AID_{50}$ *i.r.* or at $1000AID_{50}$ *i.v.* Results are shown as log p27 decrease in the presence of CD8+ T-cells. *p < 0.05, **p < 0.01, ***p < 0.001; Mann-Whitney U-test. **(C)** Plasma viral loads (left panel), $CD8^+$ T-cell activation levels (middle panel) and CD8+ T-cell-mediated SIV-suppressive activity (right panel) in CyM starting cART at day 28 post-infection. Values are shown at baseline (day -20) and different days post-infection. Gray areas indicate the period under cART. Individual values, median and interquartile range for 6 animals are shown.

**Figure S9. Dynamics of control and SIV-specific $CD8^+$ T-cell responses in M6 and non-M6 controllers.** (**A**) Evolution of $CD4^+$ T-cells and SIV-DNA levels in blood (upper panels) and peripheral lymph nodes (bottom panels) in M6 (green) and non-M6 (orange) SIV controllers. Median and interquartile range are shown. (**B**) TNF$\alpha$ production by SIV-specific $CD8^+$ T-cells (upper panels) and $CD8^+$ T-cell-mediated SIV-suppressive activity (bottom panels) in blood and peripheral lymph nodes over the course of infection and in bone marrow, spleen, and mesenteric lymph nodes at euthanasia in M6 (green) and non-M6 (orange) SICs. *p < 0.05; Mann-Whitney U-test.

**Figure S10. Flow cytometry gating strategy for the identification of SIV-specific $CD8^+$ T-cells by tetramer staining.** (**A**) Flow cytometry gating strategy used to analyze tetramer-binding $CD3^+$ $CD14^-$ $CD20^-$ $CD8^+$ T-cells. Representations are displayed as standard pseudocolor dot plots. (**B**) Representative example of tetramer staining in $CD8^+$ and $CD4^+$ T-cells from one of the animals of the study at different times post infection. The Fluorescence Minus One (FMO) control is depicted for reference.

**Figure S11. Flow cytometry gating strategy for the identification of $CD8^+$ T-cell subpopulations.** Flow cytometry gating strategy used to characterize $CD8^+$ T-cell subsets, based on expression of CD27, CD45RA, and CCR7. Gating strategy was established with baseline bulk $CD8^+$ T-cells. Representations are displayed as standard pseudocolor dot plots. Baseline expression levels of T-bet (upper panel) and CD127 (lower panel) were used to confirm the differentiation status of $CD8^+$ T-cell subsets. Naïve = dark gray; Central Memory= light blue; Transitional Memory = green; Effector Memory= yellow; Effector=red.

**Figure S12. MHC haplotype M6 and low-dose inoculation favor spontaneous control of SIV.** Plasma VL kinetics in M6 and non-M6 CyMs inoculated *i.r.* with the indicated doses of SIVmac251. Green, M6 $50AID_{50}$; orange, non-M6 $5AID_{50}$; blue, non-M6 $50AID_{50}$. Median and interquartile range are shown.

**Figure S13. Flow cytometry gating strategy used to analyze cytokine production via intracellular staining after peptide stimulation.** Representative example with PBMCs from an infected macaque stimulated with the pool of optimal SIV peptides. Results are depicted as standard pseudocolor dot plots.

**Supplemental Table 1.** Characteristics of Cynomolgus macaques included in the study.

| Monkey ID | Outcome at euthanasia | Experimental group | Age at SIV infection (years) | MHC haplotype | | |
|---|---|---|---|---|---|---|
| | | | | Class IA | Class IB | Class II |
| 29915 | SIC | M6 50AID$_{50}$ | unknown | M4/M6 | M4/M6 | M4/M6 |
| 29925 | SIC | M6 50AID$_{50}$ | unknown | M6/M2 | M6/M2 | M6/M2 |
| BA081 | SIC | M6 50AID$_{50}$ | 7.1 | M6/M1 | M6/M1 | M6/M6 |
| BA209 | SIC | M6 50AID$_{50}$ | 7.1 | M4/M6 | M4/M6 | M4/M6 |
| BC094 | SIC | Non-M6 5AID$_{50}$ | 7.4 | M3/M3 | M3M1/M3 | M1/M3 |
| BC179 | SIC | Non-M6 5AID$_{50}$ | 7.4 | M3/M1 | M3/M1 | M3/M1 |
| BC657 | SIC | M6 50AID$_{50}$ | 6.6 | M6/M2 | M6/M2 | M6/M2 |
| BD536 | SIC | Non-M6 5AID$_{50}$ | 6.5 | M3/M4 | M3/M4 | M3/M4 |
| BG927 | SIC | Non-M6 50AID$_{50}$ | 6.2 | M4/M4 | M3/M4 | M3/M4 |
| BL669 | SIC | Non-M6 50AID$_{50}$ | 5.5 | M3/M1 | M3/M1 | M3/M1 |
| BO413 | SIC | Non-M6 50AID$_{50}$ | 5 | M3/M4 | M3/M4 | M3/M4 |
| BO186 | SIC | Non-M6 5AID$_{50}$ | 4.6 | M3/M1 | M3/M1 | M3/M1 |
| 31041 | VIR | M6 50AID$_{50}$ | unknown | M6/M1 | M6/M6 | M6/M6M3 |
| AV979 | VIR | Non-M6 50AID$_{50}$ | 8.1 | M1/M5 | M1/M5 | M1/M5 |
| BB598 | VIR | Non-M6 50AID$_{50}$ | 7.1 | M3/M3 | M3/M3 | M1/M3 |
| BD885 | VIR | Non-M6 50AID$_{50}$ | 6.8 | M4/M2 | M4/M2 | M3/M2 |

**Supplemental Table 2.** List of optimal peptides used to evaluate $CD8^+$ T-cell responses.

| SIV protein | Amino acid positions | Length (amino acids) | Amino acid sequence | Primary restricting haplotype | Restricting Molecule* |
|---|---|---|---|---|---|
| Gag | 28–37 | 10 | KYMLKHVVWA | M3 | Mafa B*011:01 |
| Gag | 54–63 | 10 | KEGCQKILSV | M3 | Mafa B*075:01 |
| Gag | 146–154 | 9 | HLPLSPRTL | M3 | Mafa B*075:01 |
| Gag | 156–166 | 11 | AWVKLIEEKKF | M6 | N/A |
| Gag | 192–200 | 9 | NCVGDHQAA | M1 | Mafa B*104:01 |
| Gag | 221–229 | 9 | PAPQQGQLR | M3 | Mafa B*075:01 |
| Gag | 386–394 | 9 | GPRKPIKCW | M3 | Mafa A1*063:02 |
| Gag | 459–467 | 9 | TAPPEDPAV | M3 | Mafa B*075:01 |
| Pol | 592–600 | 9 | QVPKFHLPV | M1,M2, M3 | Mafa A4*01:01 |
| Env | 260–268 | 9 | VSSCTRMME | M3 | Mafa B*075:01 |
| Env | 338–346 | 9 | RPKQAWCWF | M3 | Mafa A1*063:02 |
| Env | 504–512 | 9 | PIGLAPTDV | M3 | Mafa B*075:01 |
| Env | 620–628 | 9 | TVPWPNASL | M3 | Mafa B*075:01 |
| Tat | 42–49 | 8 | QLYRPLEA | M3 | Mafa B1*075:01 |
| Tat | 59–67 | 9 | CCYHCQFCF | M3 | Mafa A1*063:02 |
| Rev | 26–34 | 9 | YPTGPGTAN | M3 | Mafa A1*063:02 |
| Rev | 59–68 | 10 | SFPDPPTDTP | M3 | Mafa B*075:01 |
| Nef | 103–111 | 9 | RPKVPLRTM | M3 | Mafa A1*063:02 |
| Nef | 103–112 | 10 | RPKVPLRTMS | M1,M2 | Mafa A1*063:01 |
| Nef | 194–203 | 10 | LMHPAQTSQW | M3 | Mafa B*011:01 |
| Nef | 196–203 | 8 | HPAQTSQW | M1, M2, M3 | Mafa A1*063 |
| Nef | 238–248 | 11 | GLSEEEVRRRL | M3 | Mafa B*011:01 |
| Vif | 155–163 | 9 | VVSDVRSQGE | M3 | Mafa B*011:01 |
| Vpx | 19–27 | 9 | GEAFEWLNR | M6 | Mafa B*095:01 |

*M1, M2 and M3 haplotypes transcribe strongly similar Mafa-A1*063 alleles (Budde et al., 2010). The optimal peptides described to be restricted by Mafa A1*063:01 and Mafa A1*063:02 in our table could virtually be recognized by the M1, M2 and M3 haplotypes.

**Supplemental Table 3.** Virologic and immunologic characteristics from experimental groups.

| | **Non-M6 50AID$_{50}$** | **Non-M6 5AID$_{50}$** | **M6 50AID$_{50}$** | **p** |
|---|---|---|---|---|
| % SIV controllers | 50% | 100% | 83.3% | **0.001** |
| Time to control viremia below 400 copies/mL (days *p.i.*) | 421 [105 – 421] | 47 [36 – 105] | 70 [36 – 105] | **0.041** |
| **RNA viral load** | | | | |
| Peak (Log SIV-RNA copies/mL) | 6.3 [6.0 – 6.9] | 5.9 [5.7 – 6.2] | 6.4 [5.1 – 7.1] | 0.153 |
| Time to peak (days *p.i.*) | 14 [11 – 17] | 14 [11 – 17] | 14 [11 – 17] | 1 |
| Set-point# (Log SIV-RNA copies/mL) | 4.0 [2.6 – 5.2] | 1.5 [1.1 – 1.8] | 1.3 [1.1 – 3.7] | **0.006** |
| Slope after peak viremia (1/slope peak – set-point) | $-9.7\times10^{-5}$ [$-1.3\times10^{-4}$ to $-2.3\times10^{-5}$] | $-2.7\times10^{-4}$ [$-3.2\times10^{-4}$ to $-1.2\times10^{-4}$] | $-8.7\times10^{-5}$ [$-1.5\times10^{-3}$ to $-2.1\times10^{-5}$] | 0.121 |
| **DNA viral load** | | | | |
| Peak (Log SIV-DNA copies/million CD4) | 4.3 [3.7 – 5.3] | 3.9 [3.5 –4.7] | 4.5 [3.7 – 4.7] | 0.630 |
| Time to peak (days *p.i.*) | 15 [15 – 36] | 21.5 [15 – 36] | 15 [15 – 36] | 0.769 |
| Set-point# (Log SIV-DNA copies/million CD4) | 3.7 [3.3 – 4.5] | 2.7 [2.1 – 3.0] | 2.7 [2.2 – 3.9] | **0.012** |
| Descending slope (1/slope peak – set-point) | $-9.7\times10^{-3}$ [$-4.8\times10^{-2}$ to $-4.4\times10^{-4}$] | $-1.4\times10^{-2}$ [$-3.7\times10^{-2}$ to $-2.2\times10^{-3}$] | $-4.0\times10^{-3}$ [$-3.1\times10^{-2}$ to $-1.0\times10^{-3}$] | 0.538 |
| **$CD4^+$ T-cell counts** | | | | |
| Nadir $CD4^+$ T-cells (cells/µL blood) | 241 [73 – 508] | 452 [106 – 910] | 186 [71 – 276] | 0.207 |
| Time to nadir $CD4^+$ T-cells (days) | 17 [15 – 21] | 9 [9 – 15] | 15 [9 – 36] | **0.013** |
| Set-point# $CD4^+$ T-cells (cells/µL blood) | 337 [80 – 796] | 658 [544 – 1468] | 337 [257 – 706] | 0.248 |

Median and range are indicated. p, Kruskal-Wallis H-test. # Set point defined as Month 3 post-infection

### Figure S1

## A

**Broncho alveolar lavages**

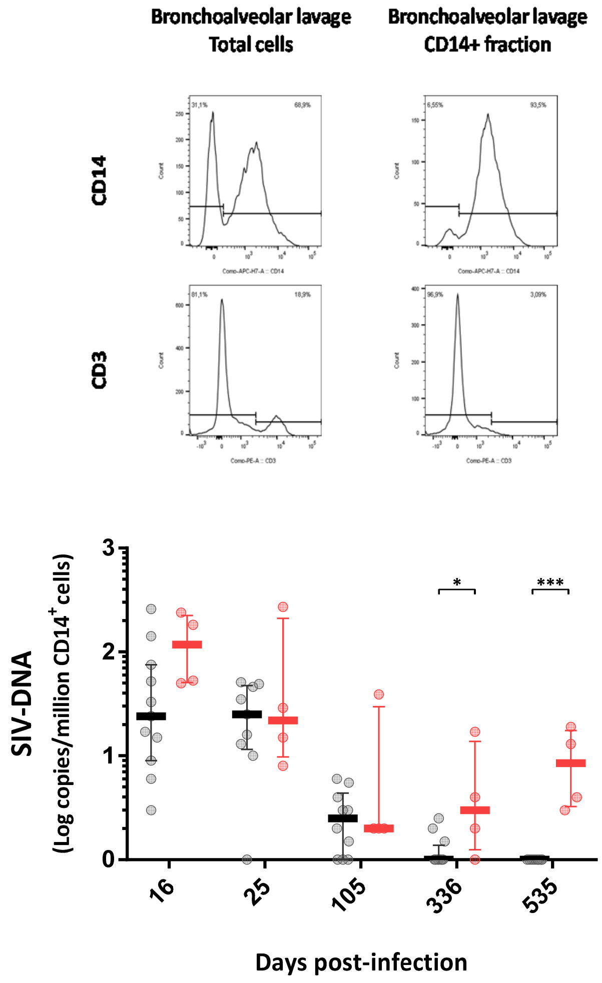


## B

**Other tissues at Euthanasia**

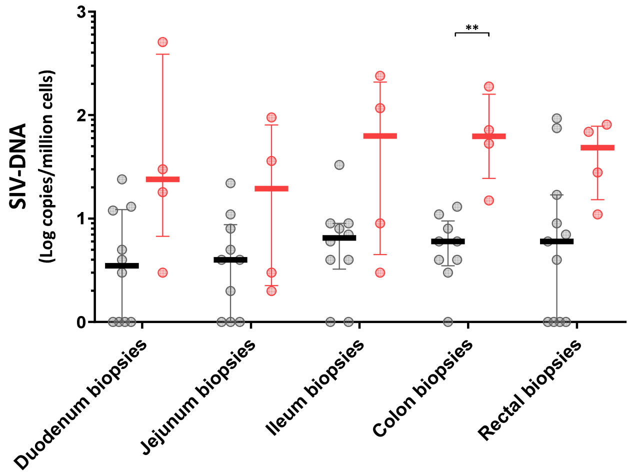


— Controllers (SICs)

— Viremics (VIRs)

#### Figure S2

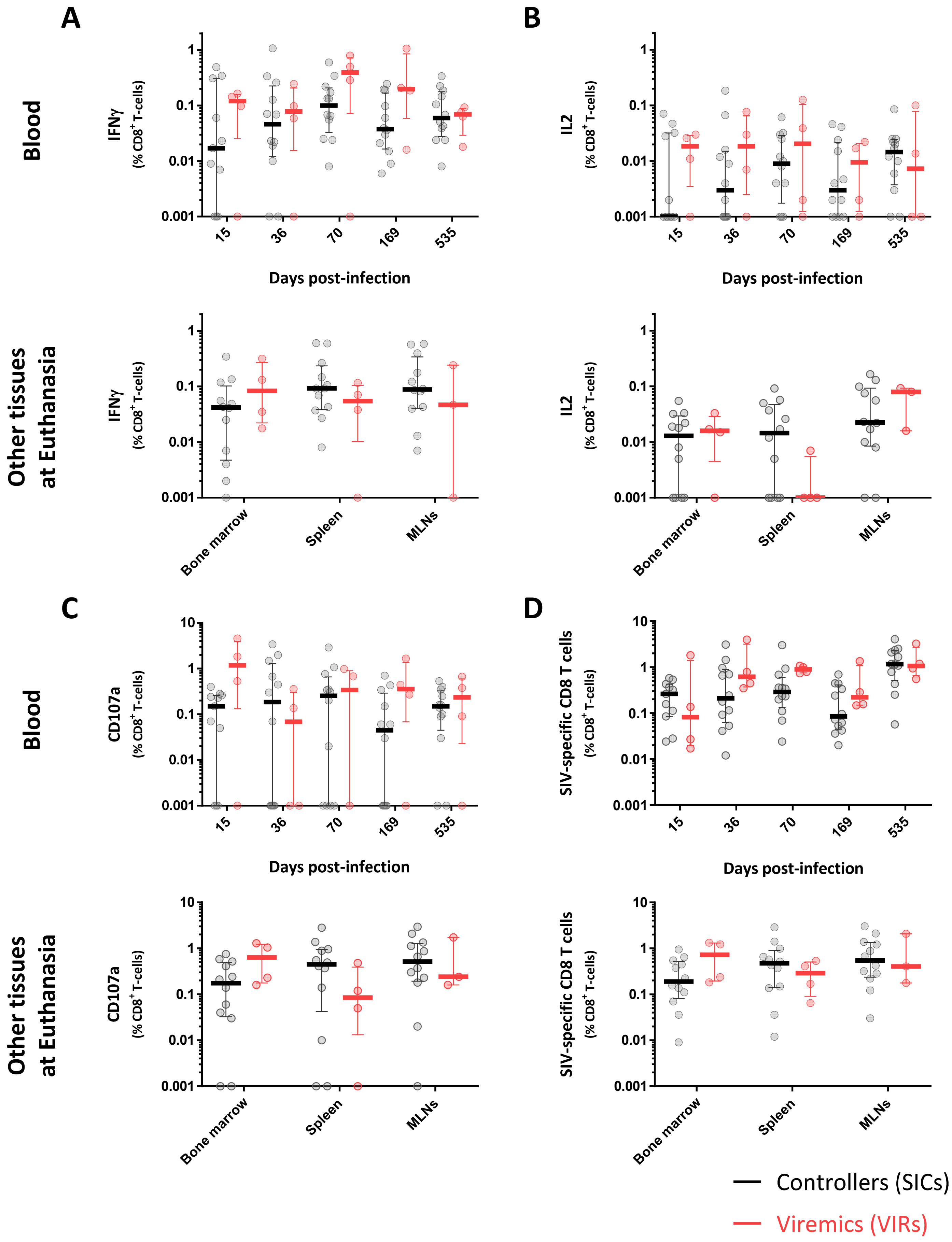

A
B
C
D
Blood
Other tissues at Euthanasia
IFNγ (% CD8+ T-cells)
IL2 (% CD8+ T-cells)
CD107a (% CD8+ T-cells)
SIV-specific CD8 T cells (% CD8+ T-cells)
Days post-infection
15
36
70
169
535
Bone marrow
Spleen
MLNs
10
1
0.1
0.01
0.001
Controllers (SICs)
Viremics (VIRs)

### Figure S3

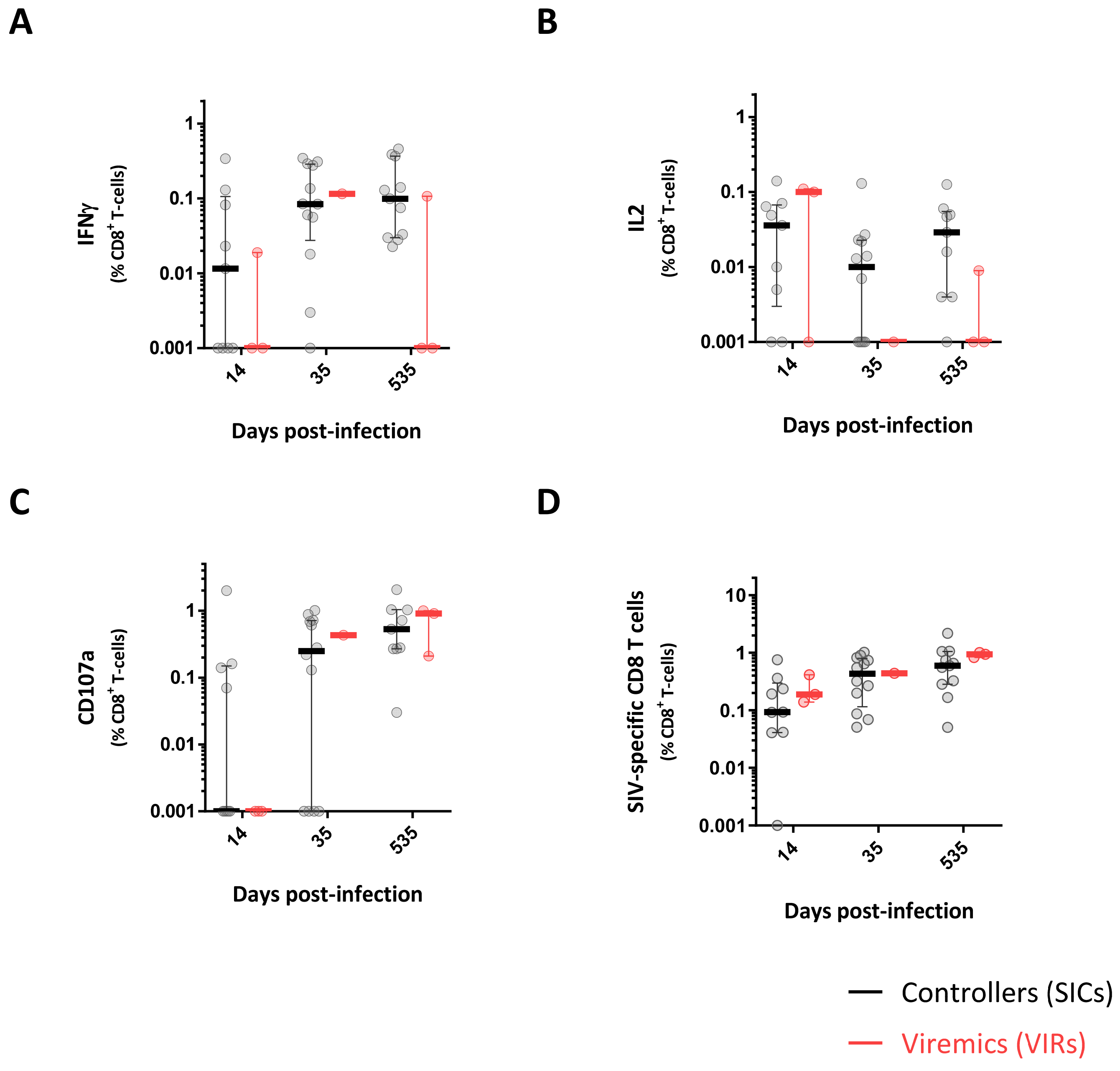
A
B
C
D
IFNγ
(% CD8+ T-cells)
IL2
(% CD8+ T-cells)
CD107a
(% CD8+ T-cells)
SIV-specific CD8 T cells
(% CD8+ T-cells)
10
1
0.1
0.01
0.001
14
35
535
Days post-infection
Controllers (SICs)
Viremics (VIRs)

### Figure S4

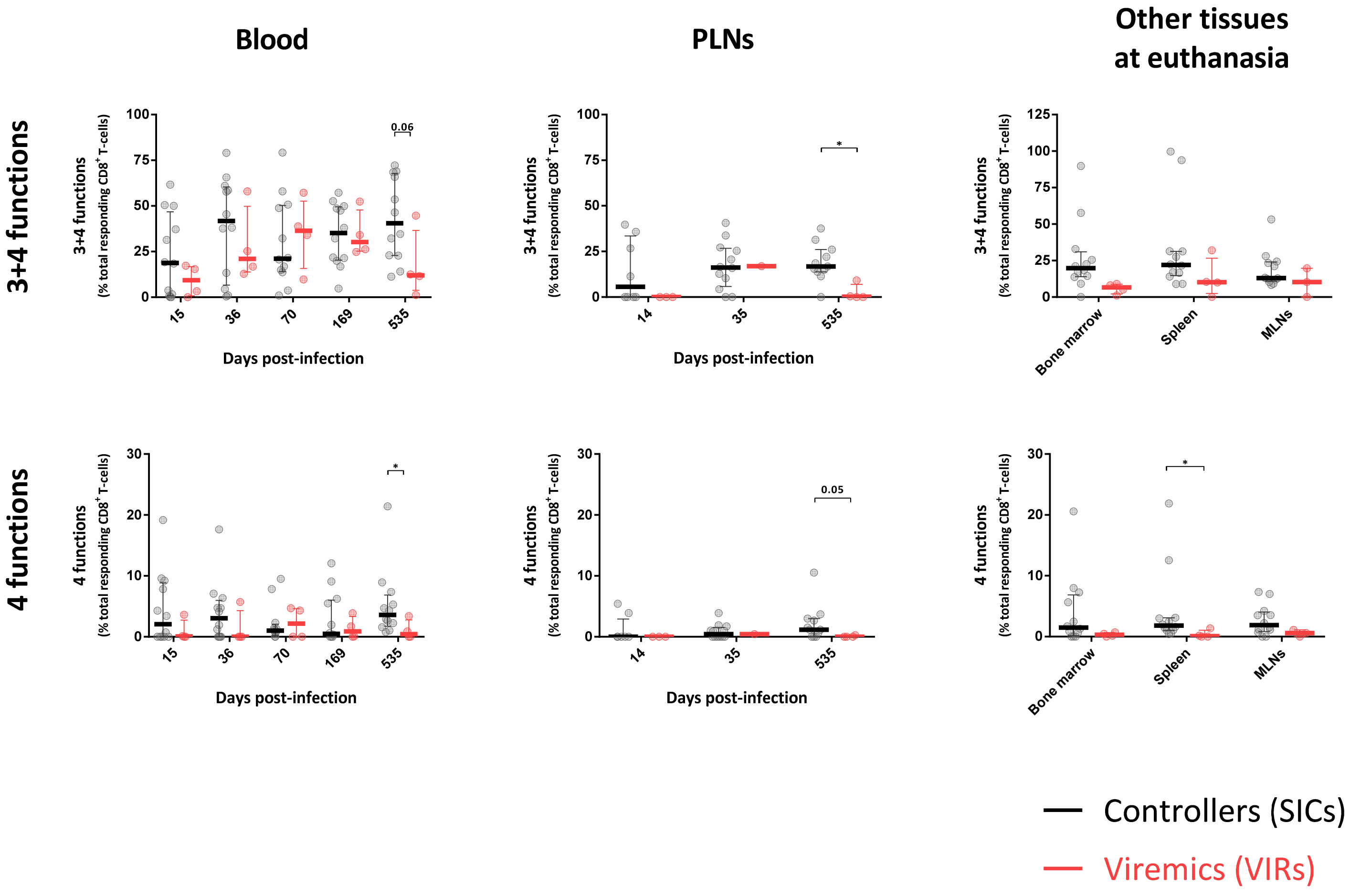
Blood
PLNs
Other tissues at euthanasia
3+4 functions
4 functions
3+4 functions (% total responding CD8+ T-cells)
4 functions (% total responding CD8+ T-cells)
Days post-infection
15
36
70
169
535
14
35
Bone marrow
Spleen
MLNs
0.06
0.05
*
Controllers (SICs)
Viremics (VIRs)

### Figure S5

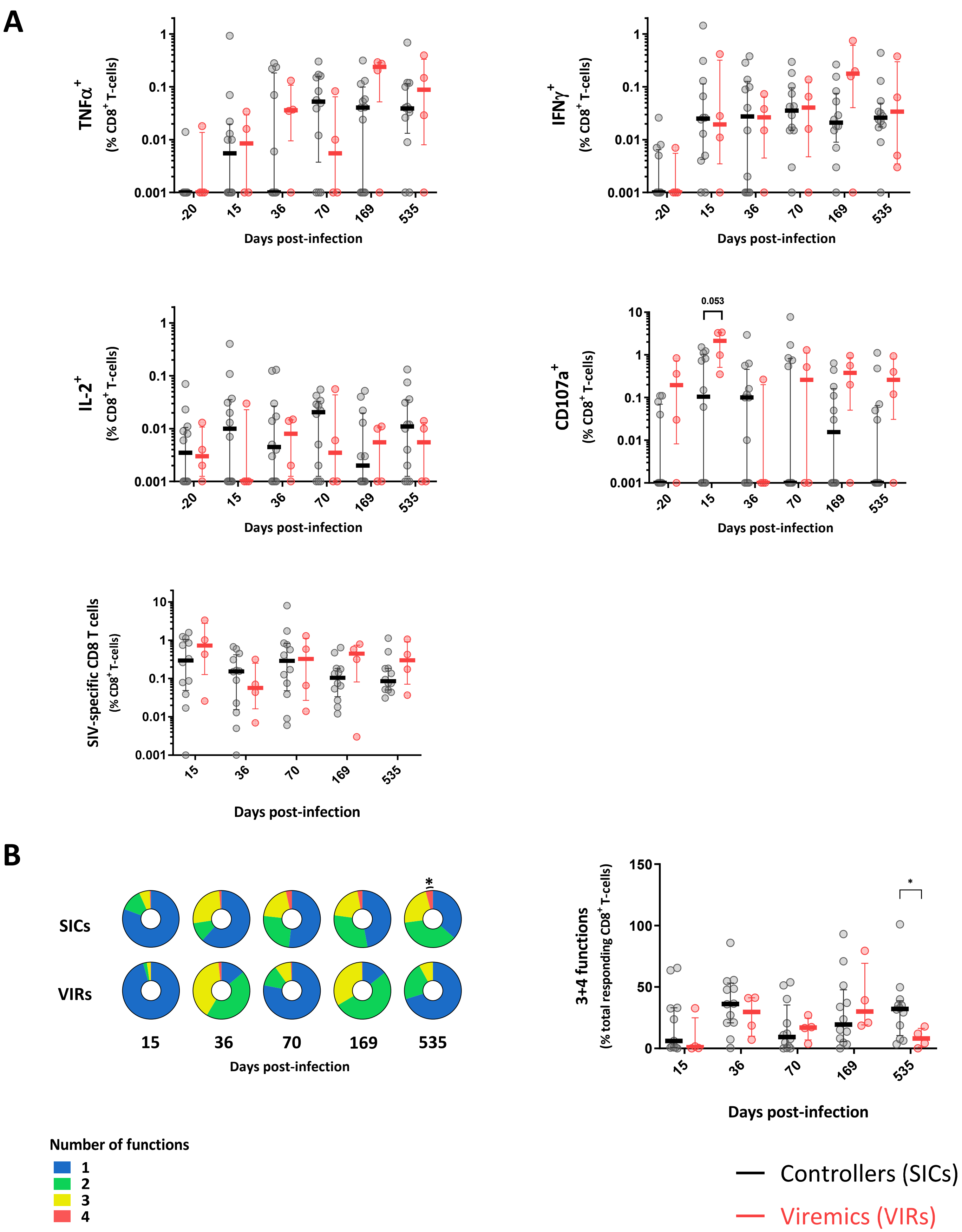

A
TNFα+ (% CD8+ T-cells)
1
0.1
0.01
0.001
-20
15
36
70
169
535
Days post-infection
IFNγ+ (% CD8+ T-cells)
IL-2+ (% CD8+ T-cells)
CD107a+ (% CD8+ T-cells)
10
0.053
SIV-specific CD8 T cells (% CD8+ T-cells)
B
SICs
VIRs
3+4 functions (% total responding CD8+ T-cells)
150
100
50
0
*
Number of functions
1
2
3
4
Controllers (SICs)
Viremics (VIRs)

#### Figure S6

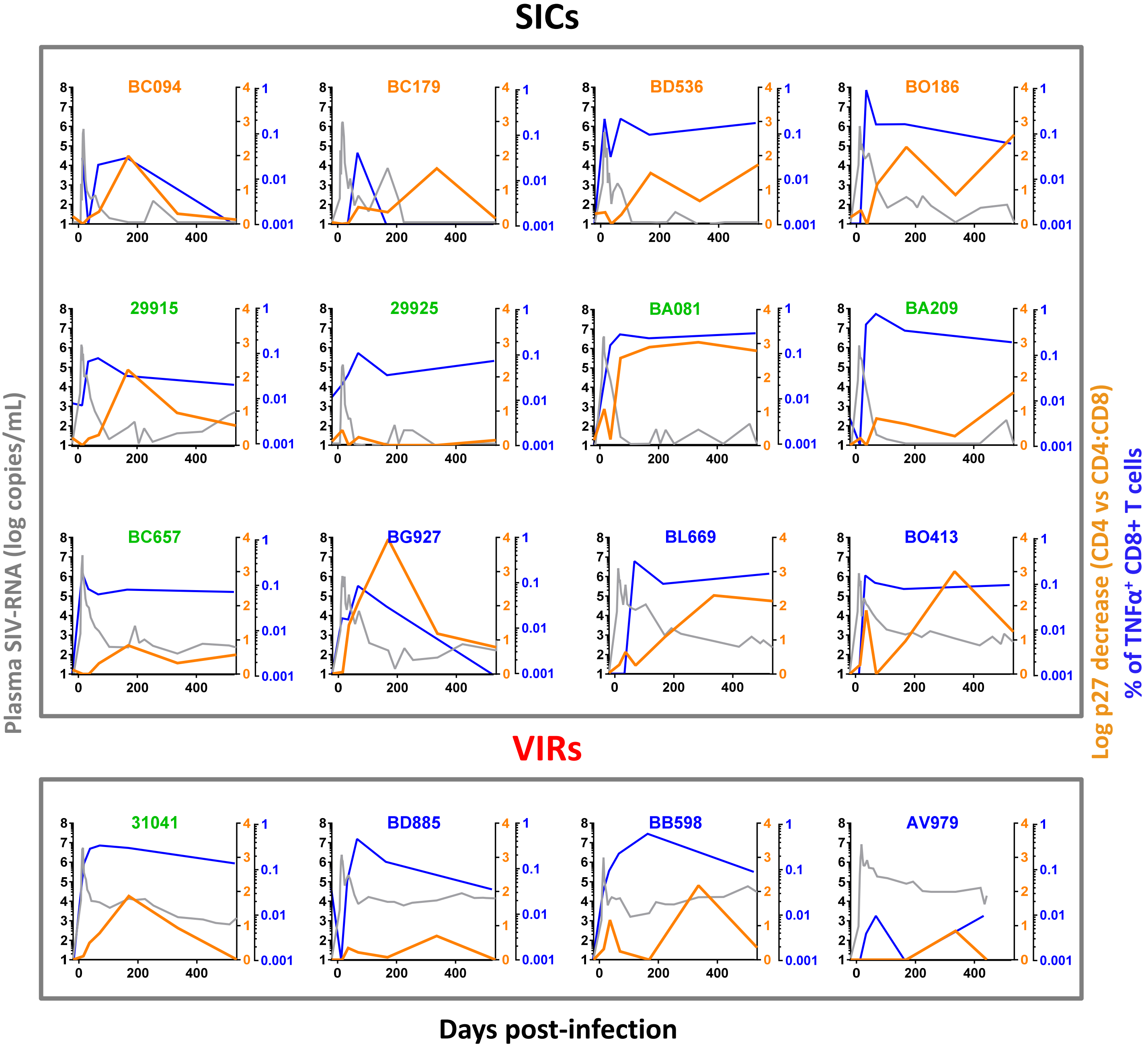

SICs
BC094
BC179
BD536
BO186
29915
29925
BA081
BA209
BC657
BG927
BL669
BO413
VIRs
31041
BD885
BB598
AV979
Plasma SIV-RNA (log copies/mL)
Log p27 decrease (CD4 vs CD4:CD8)
% of TNFα+ CD8+ T cells
Days post-infection

### Figure S7

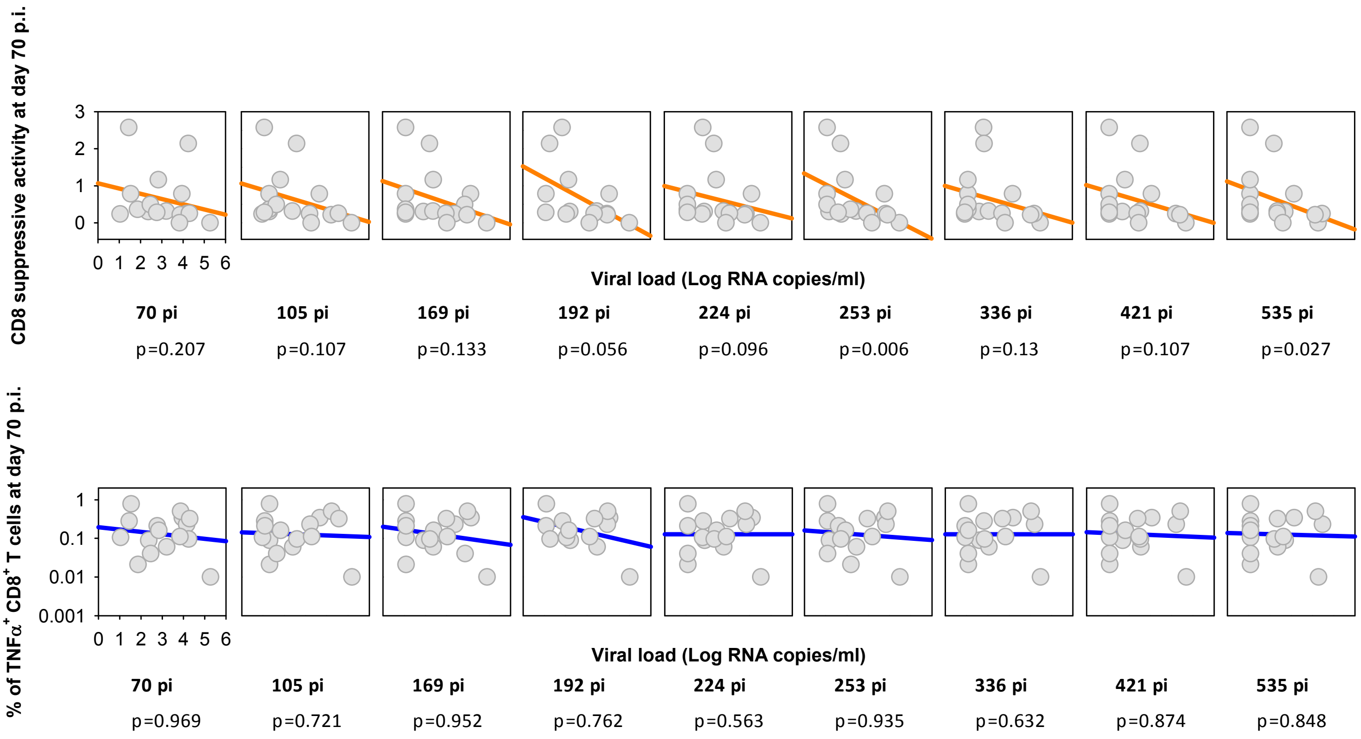

CD8 suppressive activity at day 70 p.i.
3
2
1
0
0 1 2 3 4 5 6
Viral load (Log RNA copies/ml)
70 pi
105 pi
169 pi
192 pi
224 pi
253 pi
336 pi
421 pi
535 pi
p=0.207
p=0.107
p=0.133
p=0.056
p=0.096
p=0.006
p=0.13
p=0.107
p=0.027
% of TNFα+ CD8+ T cells at day 70 p.i.
1
0.1
0.01
0.001
0 1 2 3 4 5 6
Viral load (Log RNA copies/ml)
70 pi
105 pi
169 pi
192 pi
224 pi
253 pi
336 pi
421 pi
535 pi
p=0.969
p=0.721
p=0.952
p=0.762
p=0.563
p=0.935
p=0.632
p=0.874
p=0.848

### Figure S8

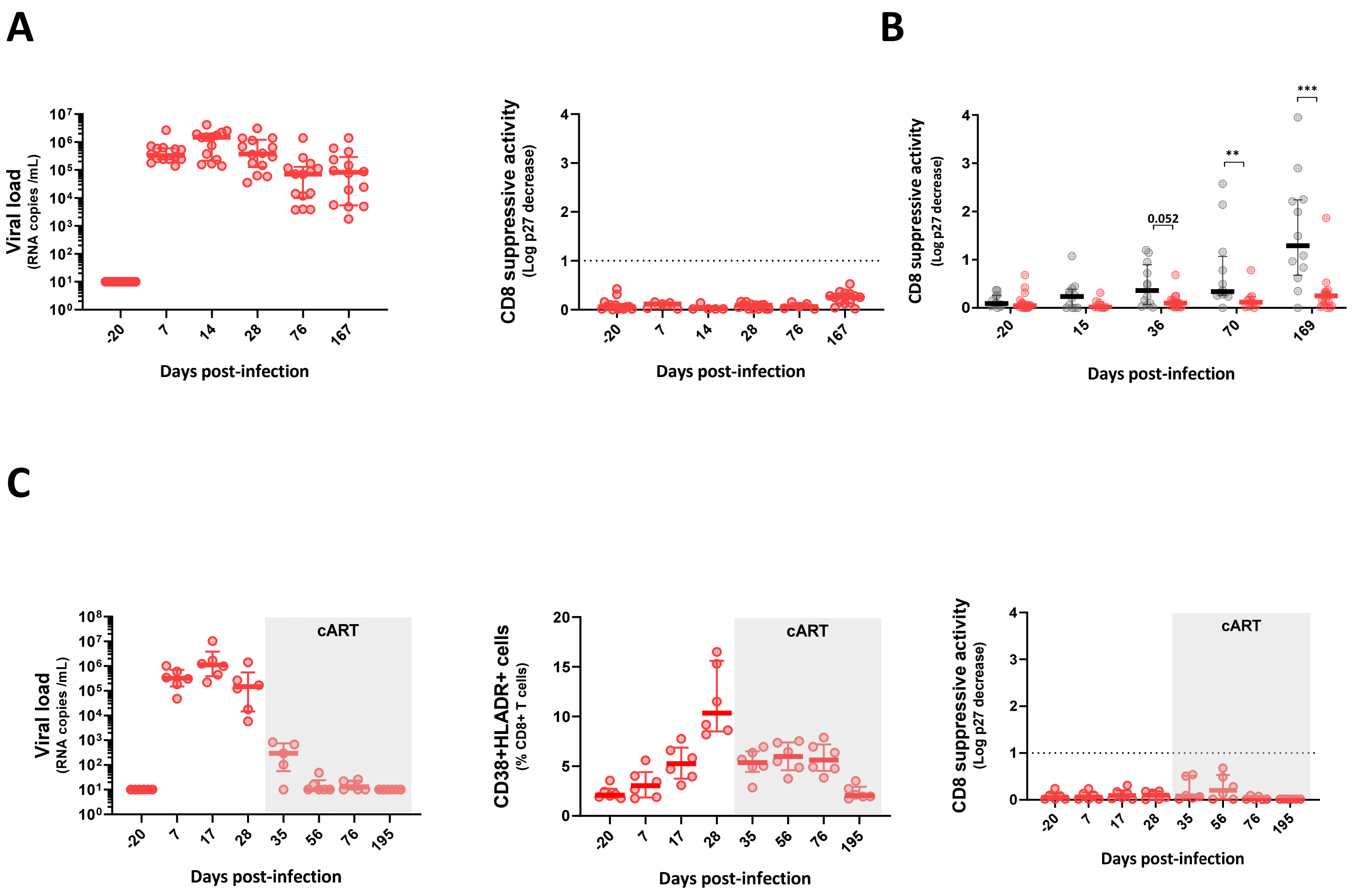
A
B
C
Viral load
(RNA copies /mL)
Days post-infection
CD8 suppressive activity
(Log p27 decrease)
0.052
**
***
cART
CD38+HLADR+ cells
(% CD8+ T cells)

### Figure S9

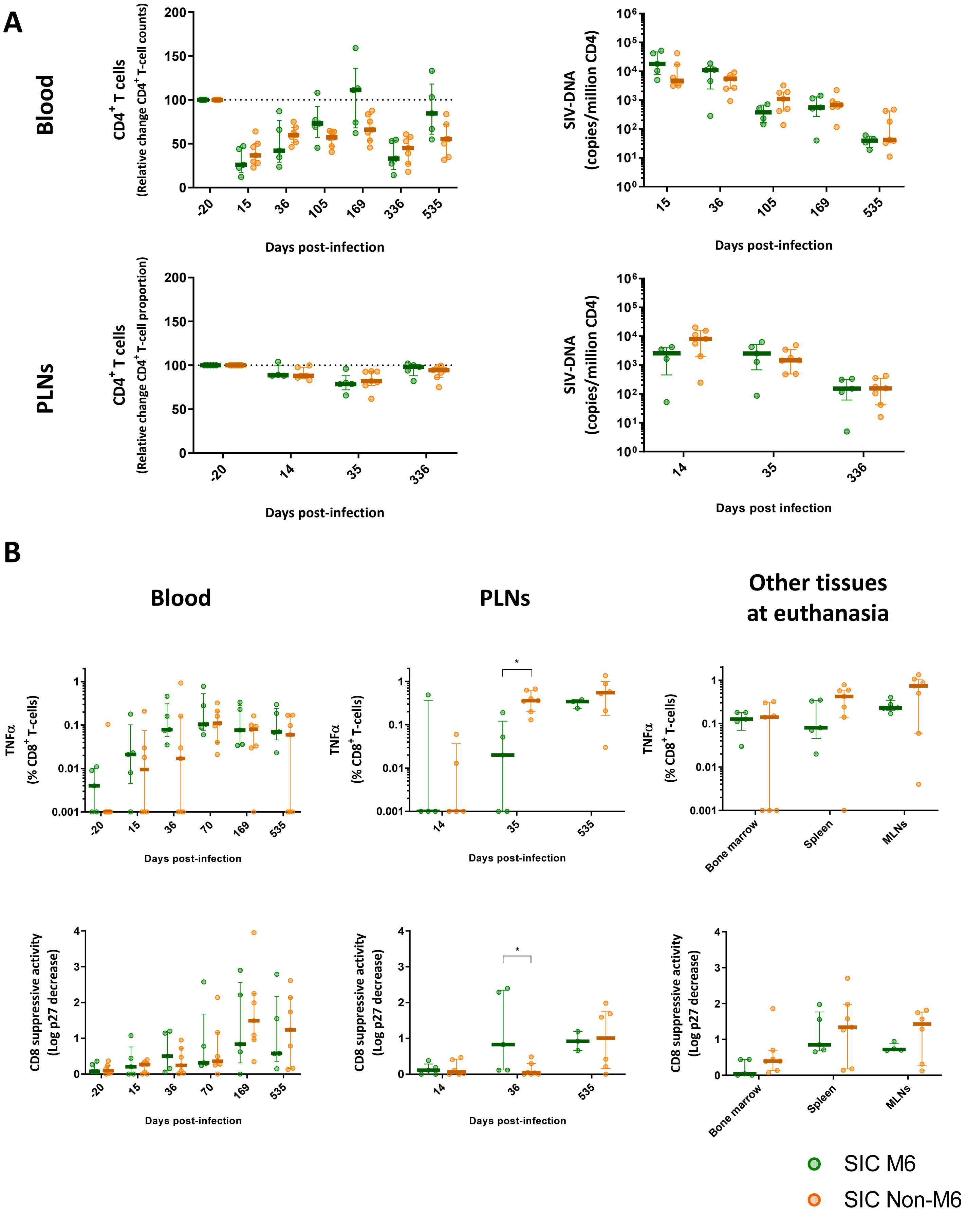
A
Blood
CD4+ T cells
(Relative change CD4+ T-cell counts)
Days post-infection
SIV-DNA
(copies/million CD4)
Days post-infection
PLNs
CD4+ T cells
(Relative change CD4+ T-cell proportion)
Days post-infection
SIV-DNA
(copies/million CD4)
Days post infection
B
Blood
PLNs
Other tissues at euthanasia
TNFα (% CD8+ T-cells)
Days post-infection
Bone marrow
Spleen
MLNs
CD8 suppressive activity (Log p27 decrease)
SIC M6
SIC Non-M6

### Figure S10

A

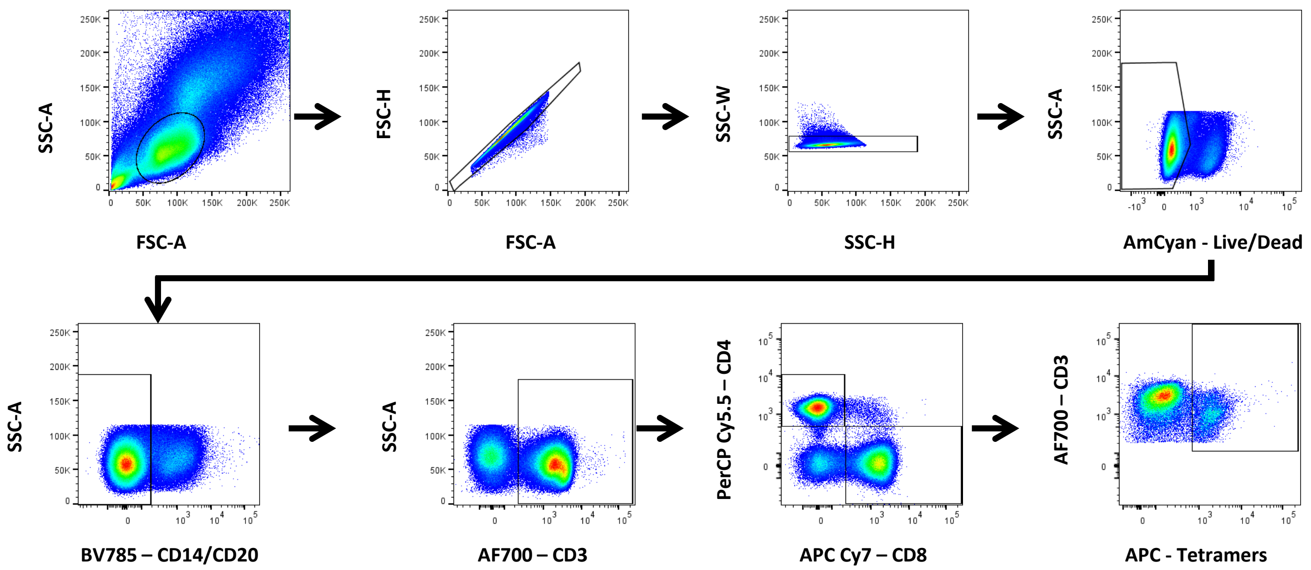


B

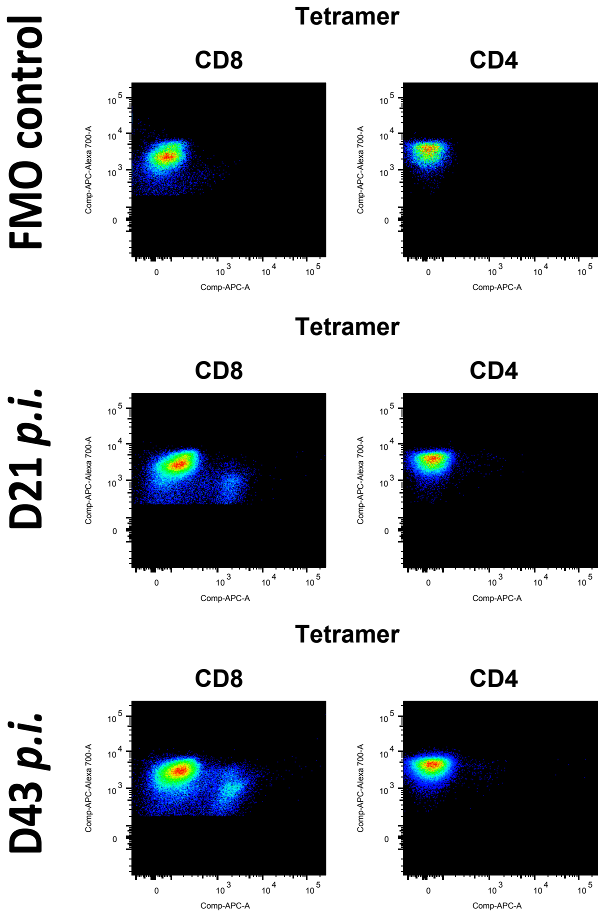

### Figure S11

## A

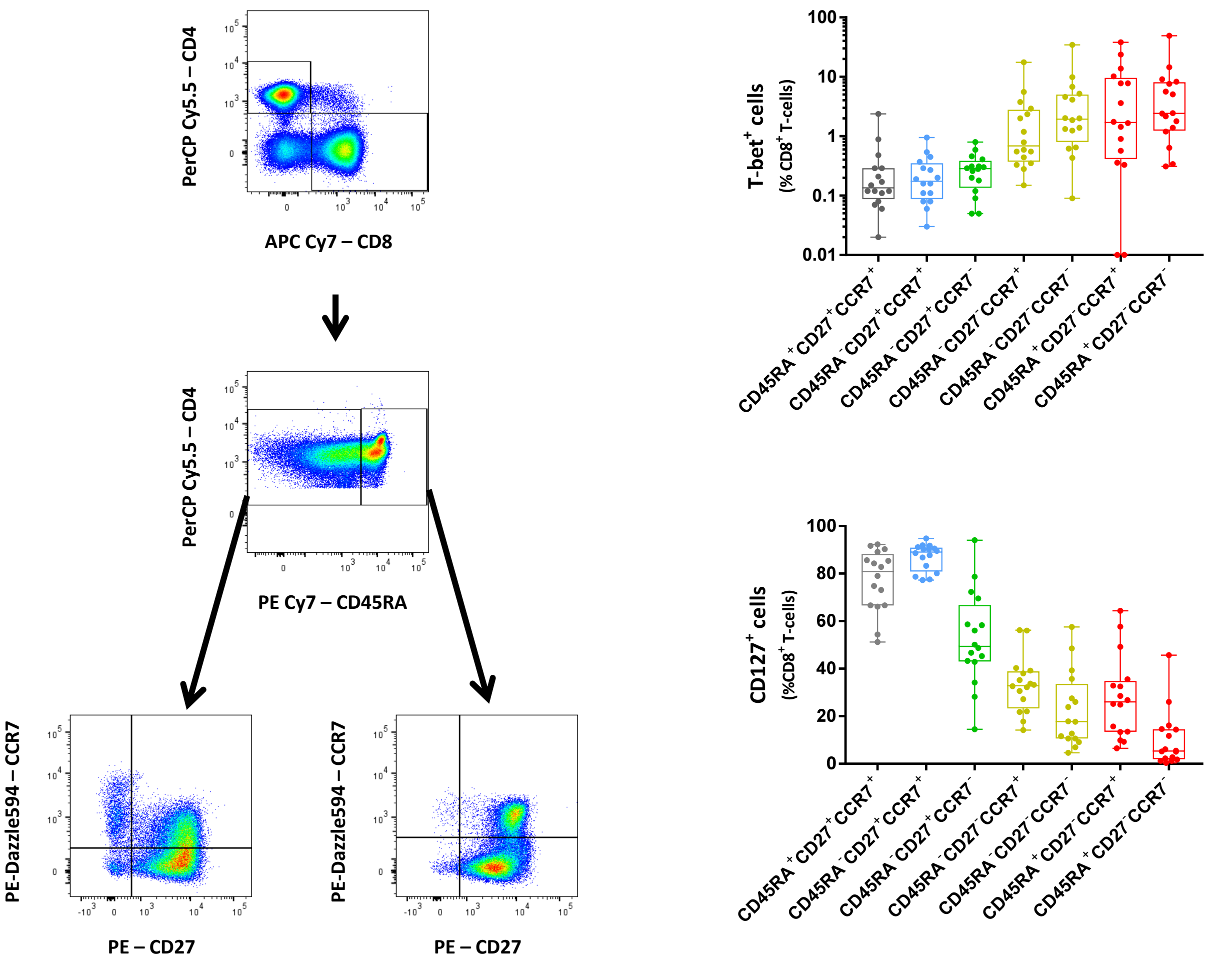
PerCP Cy5.5 – CD4
APC Cy7 – CD8
PerCP Cy5.5 – CD4
PE Cy7 – CD45RA
PE-Dazzle594 – CCR7
PE – CD27
PE-Dazzle594 – CCR7
PE – CD27
T-bet+ cells
(% CD8+ T-cells)
CD45RA+CD27+CCR7+
CD45RA-CD27+CCR7+
CD45RA-CD27+CCR7-
CD45RA-CD27-CCR7+
CD45RA-CD27-CCR7-
CD45RA+CD27-CCR7+
CD45RA+CD27-CCR7-
CD127+ cells
(%CD8+ T-cells)
CD45RA+CD27+CCR7+
CD45RA-CD27+CCR7+
CD45RA-CD27+CCR7-
CD45RA-CD27-CCR7+
CD45RA-CD27-CCR7-
CD45RA+CD27-CCR7+
CD45RA+CD27-CCR7-

#### Figure S12

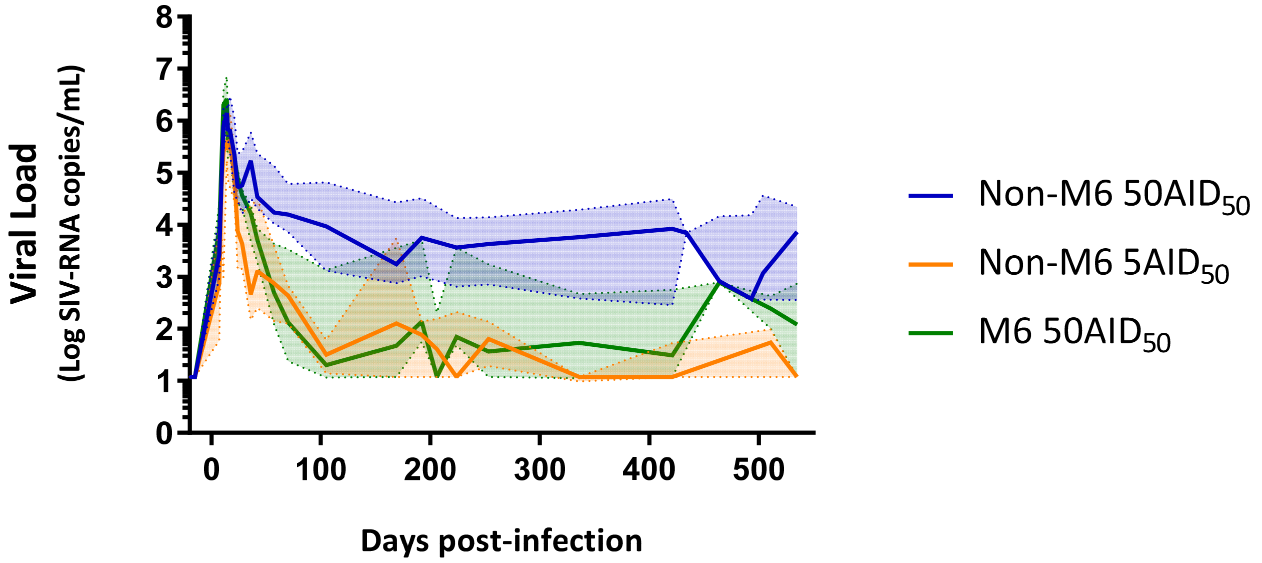

Viral Load
(Log SIV-RNA copies/mL)
8
7
6
5
4
3
2
1
0
0
100
200
300
400
500
Days post-infection
Non-M6 50AID$_{50}$
Non-M6 5AID$_{50}$
M6 50AID$_{50}$

### Figure S13

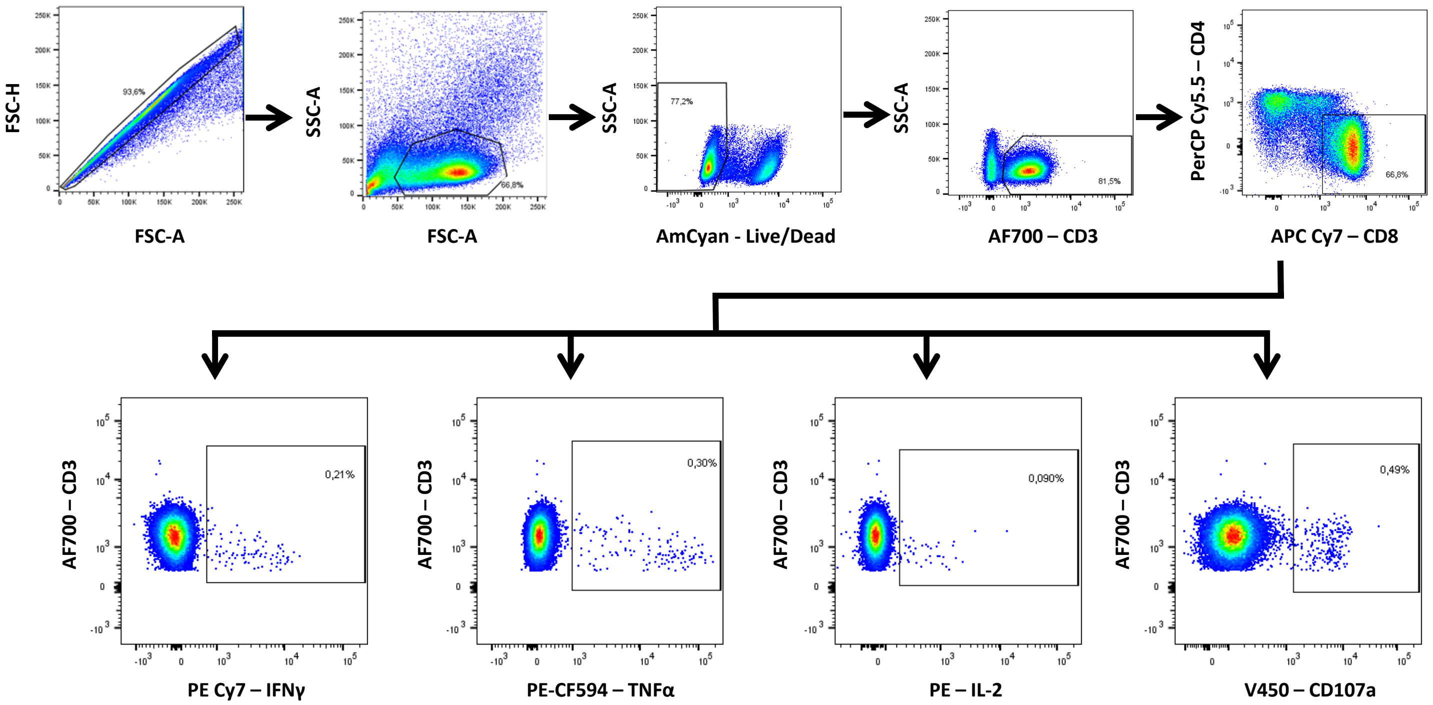

FSC-H
FSC-A
93,6%
SSC-A
FSC-A
66,8%
SSC-A
AmCyan - Live/Dead
77,2%
SSC-A
AF700 – CD3
81,5%
PerCP Cy5.5 – CD4
APC Cy7 – CD8
66,8%
AF700 – CD3
PE Cy7 – IFNγ
0,21%
AF700 – CD3
PE-CF594 – TNFα
0,30%
AF700 – CD3
PE – IL-2
0,090%
AF700 – CD3
V450 – CD107a
0,49%
